# Loss of the H4 lysine methyltransferase KMT5B drives tumorigenic phenotypes by depleting H3K27me3 at loci otherwise retained in H3K27M mutant DIPG cells

**DOI:** 10.1101/2021.07.19.452493

**Authors:** Ketty Kessler, Alan Mackay, Yura Grabovska, Valeria Molinari, Anna Burford, Sara Temelso, Haider Tari, Erika Yara-Romero, Ilon Liu, Lu Yu, David Castel, Jyoti Choudhary, Andrea Sottoriva, Mariella Filbin, Mara Vinci, Chris Jones

**Author notes:** These authors contributed equally (Ketty Kessler, Alan Mackay). Lead contact and correspondence to:* Chris Jones, Division of Molecular Pathology, The Institute of Cancer Research, 15 Cotswold Road, Sutton, Surrey, SM2 5NG, UK.

## Abstract

DIPG are characterised by histone H3K27M mutations, resulting in global loss of the repressive mark H3K27me3, although certain key loci are retained. We recently identified subclonal loss-of-function mutations in the H4 lysine methyltransferase *KMT5B* to be associated with enhanced invasion/migration, but the mechanism by which this occurred was unclear. Here we use integrated ChIP-seq, ATAC-seq and RNA-seq on patient-derived, subclonal and CRISPR-Cas9-KD DIPG cells to show that loss of *KMT5B/C* causes depletion of these retained H3K27me3 loci via changes in chromatin accessibility, causing a raft of transcriptional changes which promote tumorigenesis. De-repression occurred at bivalent loci marked by H3K4me3, driving increased transcriptional heterogeneity and elevated gene expression associated with increased invasion, abrogated DNA repair and mesenchymal transition, along with a markedly altered secretome. These data suggest a previously unrecognised trans-histone (H4/H3) interaction in DIPG cells with a potentially profound effect on their diffusely infiltrating phenotype.

## INTRODUCTION

Diffuse midline glioma with H3K27M mutation (DMG) comprise diffuse intrinsic pontine glioma (DIPG, arising in the pons) and high grade glioma occurring in other midline regions such as the thalamus (Jones and Baker, 2014). They present predominantly in young children (ages 5-10), and have a median survival of 9-12 months, with virtually all patients succumbing to the disease within 2-3 years of diagnosis (Jones et al., 2012). Radiation provides a transient response, however chemotherapeutics show little effect, and to-date targeted therapies have not provided any significant survival advantage (Jones et al., 2016). They are biologically distinct from histologically similar tumours arising in adults, harboring a key driver mutation, K27M, in genes encoding either the canonical histone H3.1 or variant H3.3 (Schwartzentruber et al., 2012; Wu et al., 2012). In a small proportion of cases, H3K27me3 is instead targeted through upregulation of the polycomb repressor complex (PRC2)-interacting protein EZHIP (Castel et al., 2020).

Despite representing a small fraction of the total H3 pool, K27M or EZHIP overexpression decreases global levels of H3K27me3, suggestive of PRC2 inhibition (Bender et al., 2013; Chan et al., 2013; Lewis et al., 2013). There are, however, areas of H3K27me3 gain at numerous loci (Bender et al., 2013; Chan et al., 2013; Funato et al., 2014), consistent with SUZ12 binding, notably at unmethylated CpG islands, which are high-affinity recruitment sites for PRC2 (Harutyunyan et al., 2019). The modification is however unable to spread to form characteristic PRC2-mediated large repressive domains (Harutyunyan et al., 2019; Lee et al., 2019; Stafford et al., 2018), and among PRC2 target genes, there is upregulation of developmental factors associated with neurogenesis at bivalent promoters (Larson et al., 2019), but also repression of *CDKN2A*, indicating enhanced self-renewal and proliferation (Cordero et al., 2017; Larson et al., 2019; Pathania et al., 2017).

A major challenge in developing novel therapeutic interventions for DIPG is the extensive intra-tumoral heterogeneity present within an individual patient’s specimen (Hoffman et al., 2016; Hoffman et al., 2019; Salloum et al., 2017). We identified for the first time genotypically- and phenotypically-distinct cancer stem-like cells to be present in these tumours, and to be retained during *in vitro* and *in vivo* model propagation (Vinci et al., 2018). Notably, we discovered novel inactivating mutations in the gene *KMT5B*, an H4 lysine methyltransferase, to be present at subclonal frequencies in DIPG specimens (<1%), and derived subclonal paired models both with and without inactivating R187* mutations (Vinci et al., 2018). *KMT5B* inactivation appeared to confer an enhanced migratory and invasive phenotype both *in vitro* and *in vivo*, in addition to a differential sensitivity to DNA repair inhibitors (Vinci et al., 2018), although the precise mechanism was unclear. Critically, these subclones were found to interact to promote tumorigenic phenotypes, whereby co-culture of *KMT5B*-deficient cells with a non-motile isogenic subclonal counterpart substantially enhanced these properties, indicating the importance of dysregulation of H4 modifications in DIPG, even when it occurs in only a small proportion of cells (Vinci et al., 2018).

KMT5B (also known as SUV420H1) and KMT5C (SUV420H2) have overlapping functions catalysing the addition of methyl groups to monomethylated histone H4K20, with a preference for H4K20me2 and H4K20me3, respectively (Jorgensen et al., 2013; Schotta et al., 2008). Disruption of these marks appear to play a key role in the maintenance of genomic integrity (Bromberg et al., 2017; Jorgensen et al., 2013; Schotta et al., 2008), and loss-of-function mutations increasingly reported in other tumour types (Brohm et al., 2019). Although less well understood, a role for H4K20 methylation and invasion has also been reported in breast cancer (Yokoyama et al., 2014). How the control of H4 modifications impact those on neighbouring histones is also unclear, however has attracted a growing interest (Bhanu et al., 2019; Bo et al., 1988; Fingerman et al., 2008; Huang et al., 2015; Liu et al., 2019). *In vitro* studies in HeLa cells have shown that nucleosomes containing the histone variant H2A.Z are enriched for H4K20me2, and that regulated replication efficiency through a KMT5B-dependent axis (Long et al., 2020). In neural stem progenitor cells of the adult subventricular zone, H3K27me3 and H4K20me3 co-localise at loci responsible for cell cycle regulation (Rhodes et al., 2016).

In the present study we explored the consequences of disrupted H4K20 methylation by KMT5B/C loss-of-function in the context of H3K27me3-dysregulated DIPG cells. We show a remarkable ablation of the retained H3K27me3 loci associated with H3K27M mutations in *KMT5B/C*-deficient cells, and identify widespread changes in chromatin accessibility to drive aberrant gene expression promoting pro-tumorigenic phenotypes. Moreover, *KMT5B* loss correlates with increased transcriptional heterogeneity and confers a poorer prognosis for children with high grade glioma.

## RESULTS

### KMT5B/C loss causes redistribution of H4K20me3 in H3K27M / EZHIP-overexpressing DIPG

In order to explore the mechanisms by which KMT5B and H4K20 modifications may play a role in DIPG tumorigenesis, we utilised several patient-derived *in vitro* experimental models (Figure 1A). From H3.3K27M-mutant HSJD-DIPG-007, we used our previously published single cell-derived subclones “F10” (*KMT5B*_R187*) and “F8” (*KMT5B* wild-type) (Vinci et al., 2018). On the background of F8, we employed CRISPR/Cas9 to engineer single gene disruption of *KMT5B* (“K5B”), *KMT5C* (“K5C”) or both (“KCo”). Furthermore, we established a novel model of H3 wild-type, *EZHIP*-overexpressing DIPG cells with a clonal, heterozygous *KMT5B*_M646fs mutation (“B193”) (Castel et al., 2020) (Figure 1B).

**Figure 1.**
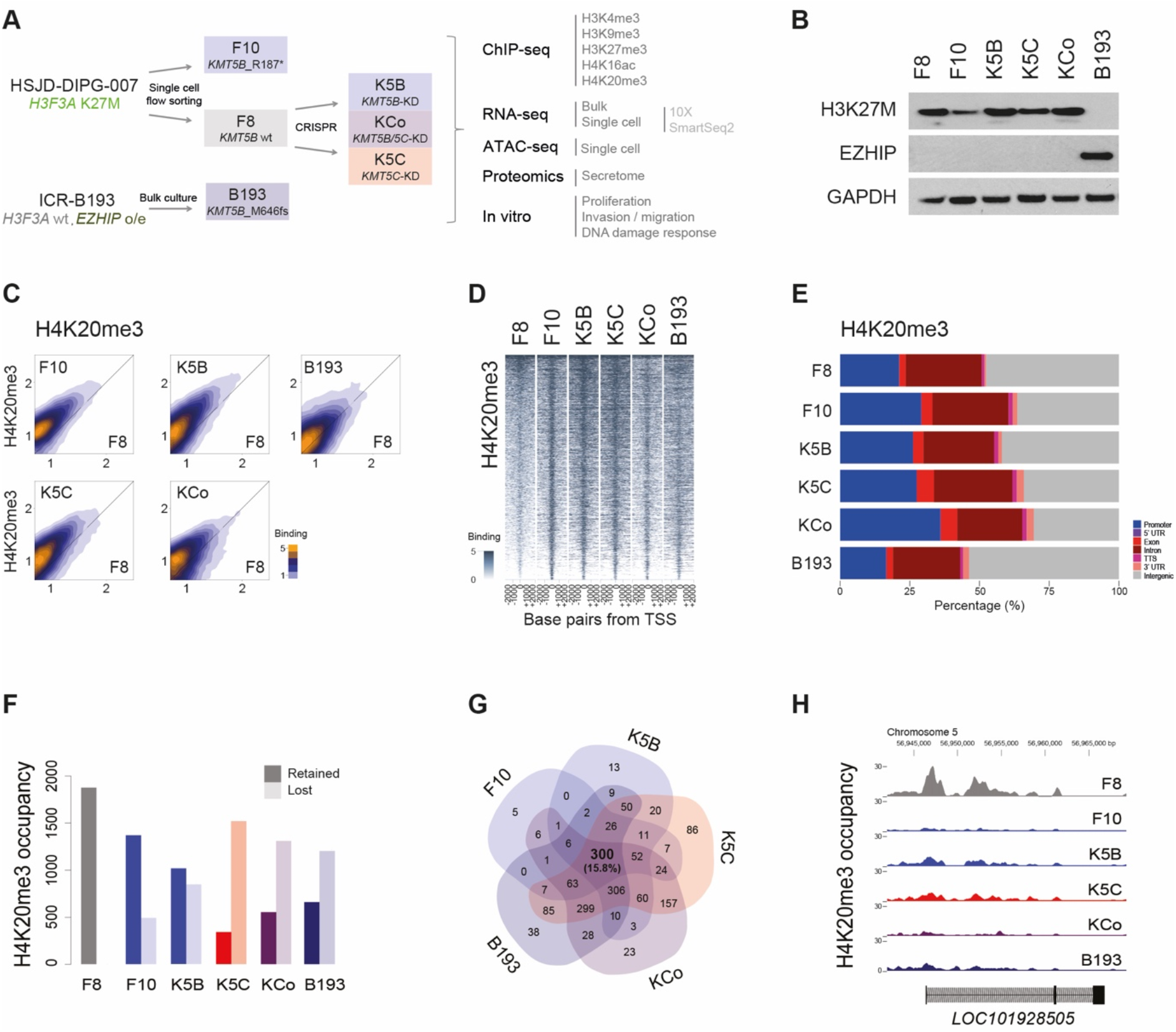
H4K20me3 binding in KMT5B/C-deficient patient-derived DIPG cells. (A) Overview of experimental design, including a subclonal flow-sorted *KMT5B*-mutant and CRISPR-KD *KMT5B/C* isogenic cells in an H3K27M background, and a clonal *KMT5*B-mutant in an *EZHIP*-overexpressing DIPG. (B) Western blot of the patient-derived models and clones used in this study, confirming expression of H3K27M or EZHIP. GAPDH is the loading control. (C) Density plot of binding of H4K20me3 peaks as assessed by ChIP-seq in *KMT5B/C-*deficient cells (y axes) compared to wild-type F8 cells (x axes). Density of peaks is coloured according to the key provided. (D) Heatmap of H4K20me3 binding in DIPG cells, ranked ordered by reads, spanning 2kb either side of the transcriptional start site. Extent of binding is coloured according to the key provided. (E) Stacked barplot showing annotated distribution of H4K20me3 binding in all cells in distinct genic and intergenic regions, coloured according to the key provided and plotted as an overall percentage per cell. (F) Barplot of H4K20me3-bound loci in F8 cells, and their retention (dark bar) or loss (light bar) in the *KMT5B/C* deficient models. (G) Venn diagram showing the overlap of H4K20me3 bound loci in F8 which are lost in the *KMT5B/C*-deficient models. (H) ChIP-seq coverage plot of H4K20me3 occupancy plotted as read depth (y axes) for each DIPG cell model for the coding sequence of *LOC101928505* on chromosome 5, with coordinates given in base pairs (x axis) and the direction of transcription from the promoter indicated. Plots from all samples are on the same scale.

Unexpectedly, however, cells with deficiencies in either *KMT5B* or *KMT5C*, but not both, showed an increase in the H4K20me2 and H4K20me3 marks that these H4 methyltransferases are responsible for depositing. This was also seen at the global level using ChIP-seq against H4K20me3, with an overall enrichment for F10, K5B, K5C and B193 cells compared to F8 cells (Figure 1C), which was predominantly associated with only low levels of occupancy across the genome (Figure 1D). The relative distributions of H4K20me3 were however shifted in the H3.3K27M cells, with an increased proportion of binding in promoters and exons relative to F8, though this was not seen in B193 (Figure 1E). This is in the context of little difference in H3K4me3 (mostly promoters) (Supplementary Figure S1A-C) or H3K9me3 (intron/intergenic) distribution (Supplementary Figure S1D-F), though there was substantially more H3K27me3 in EZHIP-overexpressing B193 compared to the H3.3K27M cells (Supplementary Figure S1G-I). There was also a reduction in H4K16ac alongside *KMT5B/C* loss (Supplementary Figure S1J-L).

Critically, genomic loci with the greatest degree of H4K20me3 occupancy in F8 were found to be lost in *KMT5B/C*-deficient cells (median 64%, range 27-81%) (Figure 1F). These were highly consistent across samples, with 15.8% loci concordant between all five cells (Figure 1G), and including intergenic as well as genic regions (Figure 1H). The intergenic regions were enriched for transcription factor binding sites such as *NRF1* (Supplementary Figure S2A) and, particularly, *TP73* (Supplementary Figure S2B). Gene ontology analysis of the genes at these loci revealed a significant enrichment of processes involved in neural development and communication (Supplementary Figure S2C). Thus, *KMT5B/C* loss redistributes H4K20me3 at important genes and regulatory regions in DIPG (Supplementary Table S1).

### Retained H3K27me3 loci in H3K27M DIPG are lost in KMT5B/C deficient cells

In addition to the changes in H4K20me3 in response to *KMT5B/C* loss, we observed that whilst there were tight correlations between all cells at loci bound by H3K4me3, there were distinct patterns in our models with other marks (Supplementary Figure S2D), both in the H3K27M (Supplementary Figure S2E) and EZHIP-overexpressing backgrounds (Supplementary Figure S2F). Most notably, there was a marked difference in F8 cells and those lacking the H4-lysine methyltransferases in respect of regions occupied by H3K27me3 (Figure 2A). There was a marked reduction in H3K27me3 at the transcriptional start sites in KMT5B/C-deficient cells (Figure 2B), far more pronounced than the changes seen with H4K20me3 (Figure 2C).

**Figure 2.**
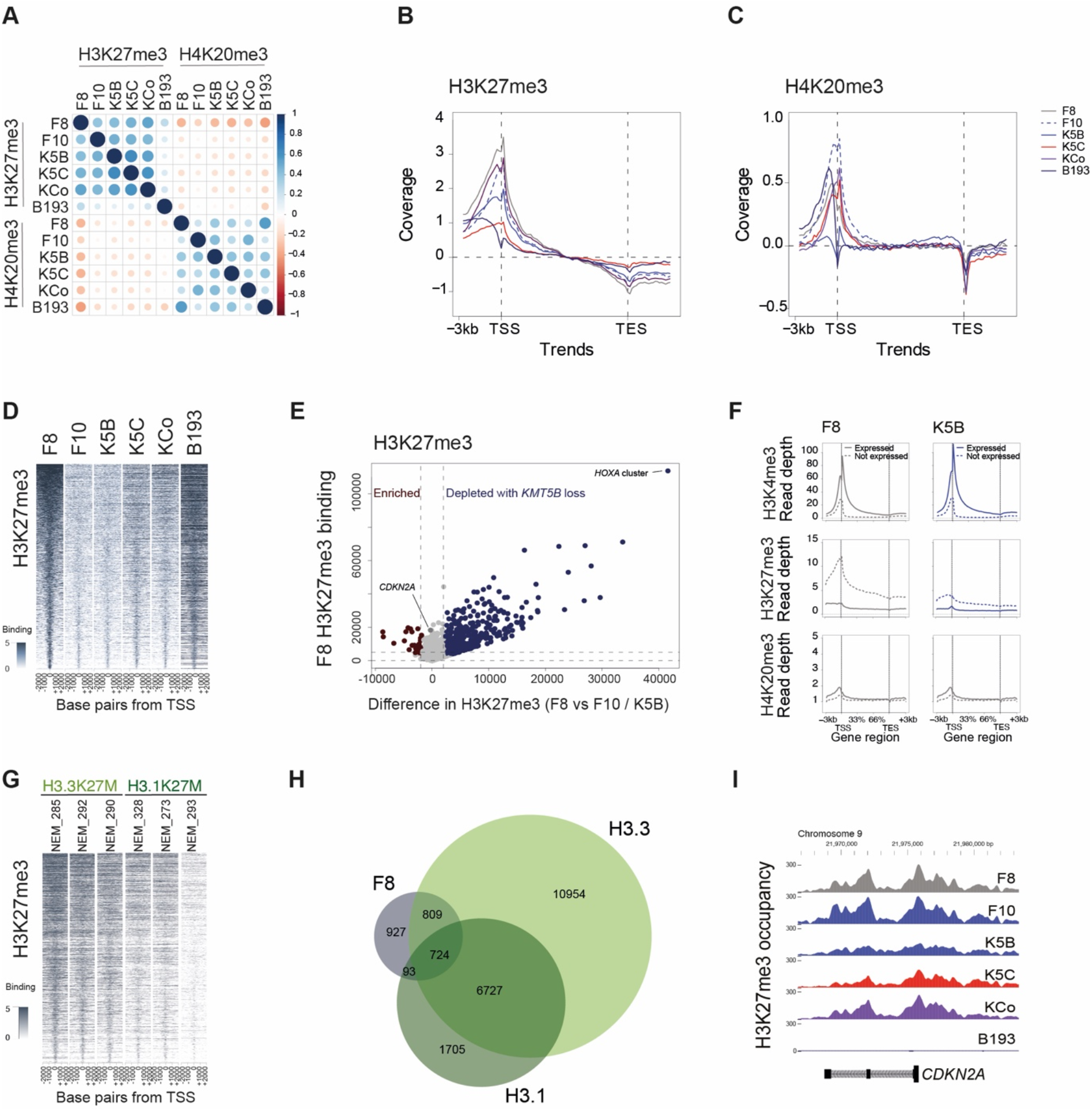
H3/H4 trans-histone effects in KMT5B-deficient DIPG cells. (A) Correlation plot of greatest H3K27me3-bound loci in *KMT5B* wild-type F8 cells against H3K27me3 and K4K20me3 loci in all cells. Size and colour of circle is dependent on the Pearson’s correlation coefficient, according to the scale provided. (B) Trend plot of coverage for H3K27me3 binding relative to +/− 3kb either side of the transcriptional start sites (TSS) and transcriptional end sites (TES) of all genes for DIPG cells as labelled. (C) Trend plot of coverage for H4K20me3 binding relative to +/− 3kb either side of the transcriptional start sites (TSS) and transcriptional end sites (TES) of all genes for DIPG cells as labelled. (D) Heatmap of H3K27me3 binding for the greatest H3K27me3-bound loci in *KMT5B* wild-type F8 in DIPG cells ranked ordered by reads, and plotted spanning 2kb either side of the transcriptional start site. Extent of binding is coloured according to the key provided. (E) Scatter of H3K27me3 binding in *KMT5B* wild-type F8 (y axis) against difference in binding between F8 and *KMT5B*-deficient F10 and K5B cells (x axis). Depleted loci with *KMT5B* loss are coloured dark blue, enriched are dark red, and no difference are grey. (F) Trend plots of coverage for H4K20me3, H3K27me3 and H3K4me3 binding relative to +/− 3kb either side of the transcriptional start sites (TSS) and transcriptional end sites (TES) of all genes for *KMT5B* wild-type F8 and -deficient K5B cells, split by genes which are expressed (solid lines) or not expressed (dashed lines). (G) Heatmap of H3K27me3 for the same loci as (D), provided for six independent DIPG samples (n=3 H3.3K27M, n=3 K3.1K27M), ranked ordered by reads, and plotted spanning 2kb either side of the transcriptional start site. Extent of binding is coloured according to the key provided. (H) Scaled Venn diagram of overlap between retained H3K27me3 loci in F8 (and subsequently lost in F10/K5B) and those retained in any of the independent H3.3K27M (light green) or H3.1K27M (dark green) samples. (I) ChIP-seq coverage plot of H3K27me3 occupancy plotted as read depth (y axes) for each DIPG cell model for the coding sequence of *CDKN2A* on chromosome 9, with coordinates given in base pairs (x axis) and the direction of transcription from the promoter indicated. Plots from all samples are on the same scale.

Although global loss of H3K27me3 is a hallmark of H3K27M mutation (Bender et al., 2013; Chan et al., 2013; Lewis et al., 2013), certain loci retain occupancy (Bender et al., 2013; Chan et al., 2013; Funato et al., 2014). We saw a remarkable ablation of these retained H3K27me3 marks seen in the *KMT5B*-deficient cells, and to a lesser degree the *KMT5C*-deficient and double-KD (Figure 2D). This was true at all levels of H3K27me3 occupancy in F8, with 2553/4641 (55%) of bound loci lost in F10 and/or K5B cells (Figure 2E). Notably, this transhistone effect on H3K27me3 had a profound effect on transcription, with the inhibitory effect on gene expression exerted by H3K27me3 in *KMT5B* wild-type F8 cells absent with *KMT5B*-KD, although H3K4me3 was unchanged (Figure 2F). To investigate whether these loci were cell line-dependent, we assessed H3K27me3 levels in these regions in further H3.3K27M (n=3) and H3.1K27M samples (n=3) (Castel et al., 2018) (Figure 2G). There was a high degree of concordance in retention of these loci in 5/6 tumours, more so in H3.3 than H3.1 K27M tumours, and with a total of 1626/2553 (64%) common with those in our models (Figure 2H) (Supplementary Table S2). Importantly, some of these ‘retained’ loci were not subsequently lost with *KMT5B/C* deficiency, and included key genes reported to play an important role in DIPG tumorigenesis such as *CDKN2A/B* (Mohammad et al., 2017) (Figure 2I), though curiously this locus was lost in B193.

### KMT5B-driven transhistone interactions drives aberrant expression of bivalent genes marked by H3K4me3

The genomic regions in which otherwise retained H3K27me3 occupancy was lost in the absence of *KMT5B/C* encompassed both relatively large loci spanning several genes, such as *GLI1* / *ARHGAP9* (Figure 3A), *NRGN* / *VSIG2* / *ESAM* (Figure 3B) as well as individual genes such as *IGFBP3* (Figure 3C), *STMN2* (Supplementary Figure S3A), *MARCH4* (Supplementary Figure S3B) and *DRD2* (Supplementary Figure S3C). This was common across the single gene mutation/KD F10, K5B and K5C cells (all p<0.0001, t-test) (Figure 3D). This reduction in H3K27me3 binding resulted in increased gene expression by RNAseq (p<0.0001, t-test) (Figure 3E). In 90/98 (92%) of these loci, gene promoters retained H3K4me3 binding in all samples (Supplementary Figure S3D), with no qualitative difference in H3K4me3 occupancy (Supplementary Figure S3E), and increased gene expression compared to the few instances lacking H3K4me3 binding (p=0.030, F10; p=0.045, K5B, t-test) (Supplementary Figure S3F). Thus, we are able to identify a common set of genes with increased expression upon *KMT5B* loss which are defined by a loss of H3K27me3 at bivalent promoters (Figure 3F) (Supplementary Table S3).

**Figure 3.**
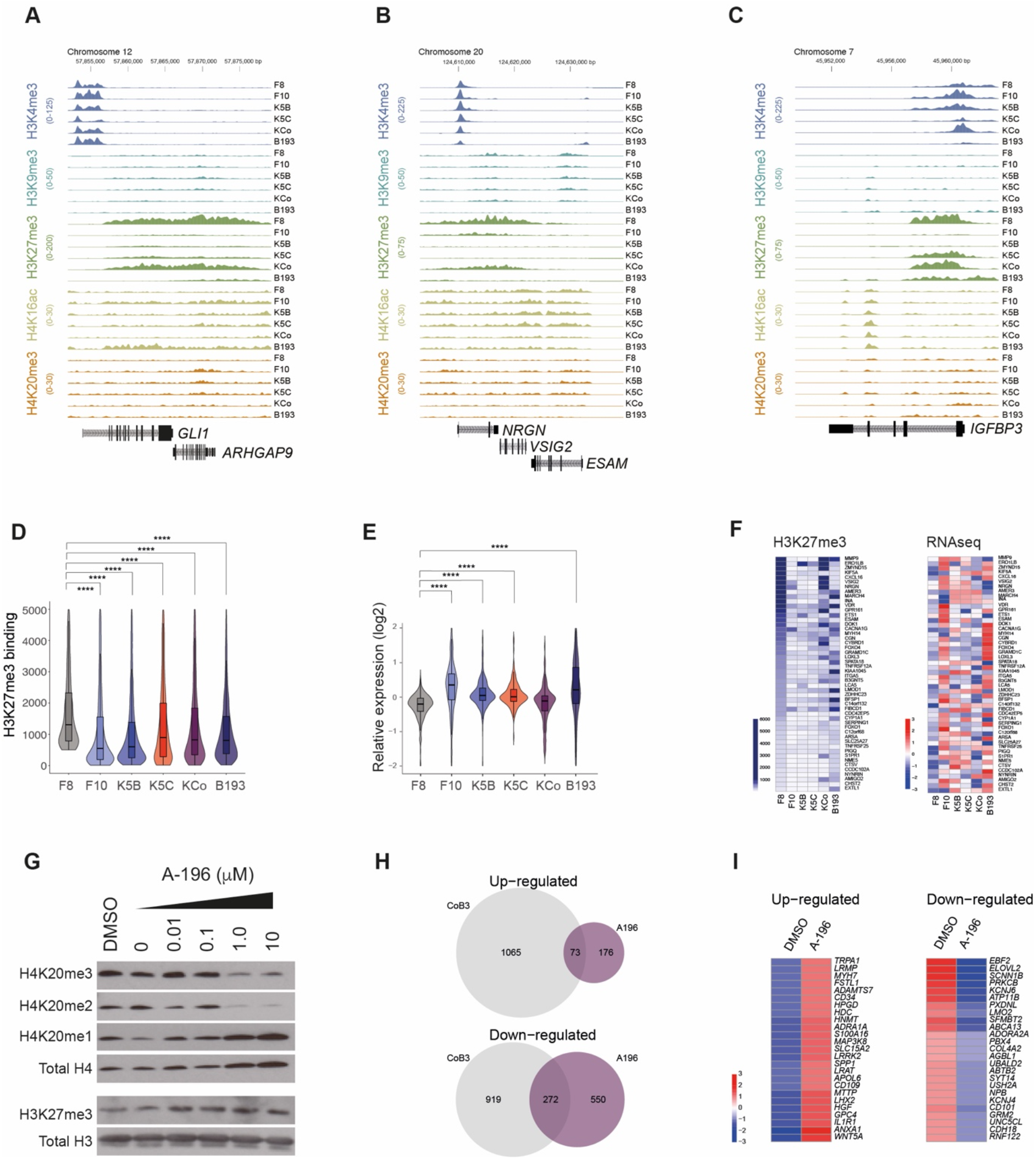
Loss of H3K27me3 binding in KMT5-deficient cells drives aberrant gene expression. (A) ChIP-seq coverage plot of H3K4me3 (blue), H3K9me3 (aqua), H3K27me3 (green), H4K16ac (mustard) and H4K20me3 (orange) occupancy plotted as read depth (y axes) for each DIPG cell model at the *GLI1* / *ARHGAP9* locus on chromosome 12, with coordinates given in base pairs (x axis) and the direction of transcription from the promoter indicated. Plots from all samples for any given mark are on the same scale. (B) ChIP-seq coverage plot of H3K4me3 (blue), H3K9me3 (aqua), H3K27me3 (green), H4K16ac (mustard) and H4K20me3 (orange) occupancy plotted as read depth (y axes) for each DIPG cell model at the *NRGN* / *VSIG2 / ESAM* locus on chromosome 20, with coordinates given in base pairs (x axis) and the direction of transcription from the promoter indicated. Plots from all samples for any given mark are on the same scale. (C) ChIP-seq coverage plot of H3K4me3 (blue), H3K9me3 (aqua), H3K27me3 (green), H4K16ac (mustard) and H4K20me3 (orange) occupancy plotted as read depth (y axes) for each DIPG cell model at the *IGFBP3* locus on chromosome 7, with coordinates given in base pairs (x axis) and the direction of transcription from the promoter indicated. Plots from all samples for any given mark are on the same scale. (D) Violin plot of H3K27me3 binding of highly bound loci in *KMT5B* wild-type, H3.3 K27M F8 cells, for F8 as well as KMT5/C-deficient cells. The horizontal line with the boxes represents the median, the box the interquartile range, and the whiskers spanning 95% of the data. *** p<0.0001, t-test. (E) Violin plot of mRNA expression for genes within the highly H3K27me3-bound loci in *KMT5B* wild-type, H3.3 K27M F8 cells, for F8 as well as *KMT5B/C*-deficient cells. The horizontal line with the boxes represents the median, the box the interquartile range, and the whiskers spanning 95% of the data*** p<0.0001, t-test. (F) Heatmap of the most differentially H3K27me3-bound and expressed genes between *KMT5B* wild-type, H3.3 K27M F8 cells and *KMT5B/C*-deficient cells. Genes are arranged by hierarchical clustering, and binding or expression represented according to the keys provided. (G) Western blot for H4K20me3, H4K20me2, H4K20me1 (with total H4 as loading control), and H3K27me3 (with total H3 as loading control) for *KMT5B* wild-type, H3.3 K27M F8 cells in response to increasing concentrations of the dual KMT5B/C inhibitor A-196. (H) Scaled Venn showing overlap in differentially expressed genes in F8 cells treated with 10μM A-196 for 24hours versus vehicle (purple) compared to CoB3 *versus* F8 (grey). (I) Heatmap of differentially expressed gene in F8 cells treated with 10μM A-196 for 24hours versus vehicle RNAseq heatmap after A-196 treatment. Expression is represented according to the keys provided.

Such an effect was also noted by pharmacological inhibition of KMT5B/C by the selective inhibitor A-196 (Bromberg et al., 2017), which causes reduction in global H4K20me2 and H4K20me3 (Figure 3G) in the absence of direct effects on enzyme protein levels (Supplementary Figure S3G) or cell viability (Supplementary Figure S3H) in F8 cells. Analysing RNAseq data on cells treated with the drug for 24hrs at 1µM, we found 345 shared differentially expressed genes in common with the double-KD cells, with a higher degree of overlap in those down-regulated (Figure 3H). Such genes (Figure 3I) were significantly enriched for process associated with cell adhesion, integrin signalling and interleukin secretion, amongst others, by gene ontology analysis (Supplementary Figure S3I). Thus, the KMT5B/C-associated trans-histone effects on gene expression appear to be directly mediated by H4K20me2/me3 deposition.

### H3/H4-mediated gene expression changes drive functionally relevant processes in DNA repair and tumour cell invasion

To explore the functional consequences of the observed epigenomic/transcriptomic changes, we first focussed on the DNA damage response, a process known to be mediated by H4K20me2/me3. The histone acetyltransferase *KAT5* (TIP60) was one the genes identified as having lost H3K27me3 in our models, (Supplementary Figure 4A), and treatment with 2Gy ionising radiation (IR) resulted in a significant reduction in the number of 53BP1 foci observed in *KMT5B*-deficient cells (p=0.0376 - <0.0001, ANOVA) (Supplementary Figure 4B). These models were also significantly more sensitive to DNA repair modulation such as the PARP inhibitor talazoparib (p=0.0209 - <0.0001, ANOVA) (Supplementary Figure 4C), as we had previously demonstrated for our single cell-derived subclones (Vinci et al., 2018). Furthermore, we observed significant enrichment of genes associated with these processes by GSEA, including DNA repair signatures (p=1.59×10^−5^) (Supplementary Figure 4D), more specifically those associated with homology-directed repair (p=3.85×10^−5^) (Supplementary Figure 4E) and cell cycle genes post-IR (p=6.43×10^−11^) (Supplementary Figure 4F).

Having established our models to recapitulate certain predicted phenotypic changes, we next explored the impact of our novel mechanism to promote tumour invasion. We observed several genes to be upregulated via H3K27me3 loss at bivalent loci, including *MMP9* (Figure 4A), *MMP24* (Figure 4B) and *ITGA5* (Figure 4C). This was underpinned by consistent enrichment of relevant gene expression signatures by GSEA, including those associated with epithelial-mesenchymal transition (p=7.26×10^−13^) (Figure 4D), extracellular matrix reorganization (p=3.15×10^−8^) (Figure 4E), and integrin pathways (p=2.96×10^−6^) (Figure 4F). Immunofluorescent staining of cells disseminating out from neurospheres and allowed to invade through Matrigel demonstrated the expected loss of H4K20me2 and/or H4K20me3 directly in the invading cell populations (Figure 4G). Despite having similar proliferation rates, loss of KMT5B specifically enhanced invasion (Figure 4H), particularly in F10 (p=2.74×10^−14^, t-test), but also in K5B, compared to F8 (p=8.72×10^−13^, t-test) (Figure 4I).

**Figure 4.**
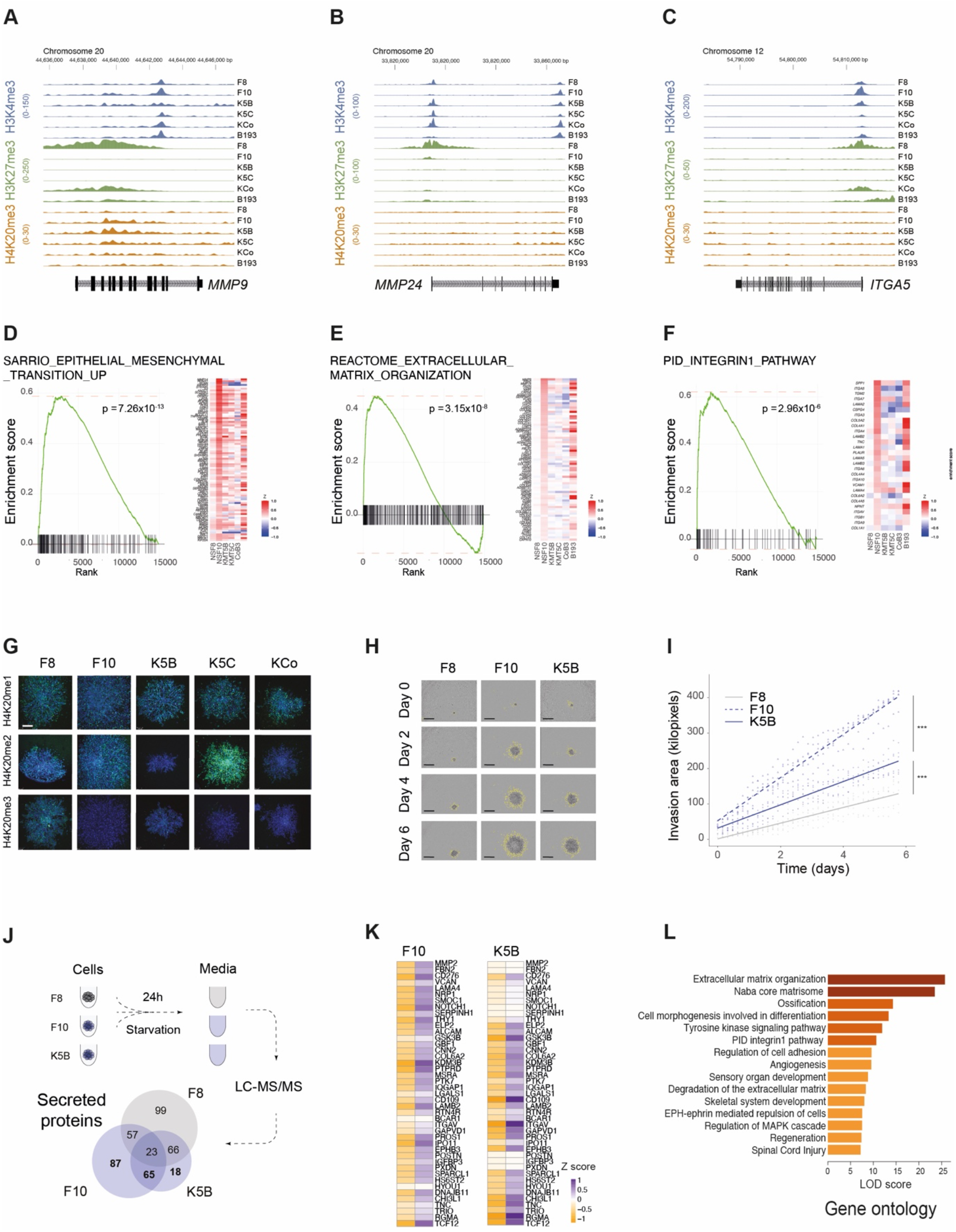
Loss of KMT5B causes gene expression changes promoting DIPG cell invasion via H3K27me3/H3K4me3. (A) ChIP-seq coverage plot of H3K4me3 (blue), H3K27me3 (green) and H4K20me3 (orange) occupancy plotted as read depth (y axes) for each DIPG cell model for the coding sequence of *MMP9* on chromosome 20, with coordinates given in base pairs (x axis) and the direction of transcription from the promoter indicated. Plots from all samples for any given mark are on the same scale. (B) ChIP-seq coverage plot of H3K4me3 (blue), H3K27me3 (green) and H4K20me3 (orange) occupancy plotted as read depth (y axes) for each DIPG cell model for the coding sequence of *MMP24* on chromosome 20, with coordinates given in base pairs (x axis) and the direction of transcription from the promoter indicated. Plots from all samples for any given mark are on the same scale. (C) ChIP-seq coverage plot of H3K4me3 (blue), H3K27me3 (green) and H4K20me3 (orange) occupancy plotted as read depth (y axes) for each DIPG cell model for the coding sequence of *ITGA5* on chromosome 12, with coordinates given in base pairs (x axis) and the direction of transcription from the promoter indicated. Plots from all samples for any given mark are on the same scale. GSEA enrichment plots for a mesenchymal transition signature in *KMT5B*-deficient DIPG cells compared to wild-type F8. The curves show the enrichment score on the y axis and the rank list metric on the x axis. Alongside is a heatmap representation of expression of significantly differentially expressed genes in the signature in all KMT5B/C-deficient DIPG cells compared to wild-type F8. (E) GSEA enrichment plots for an ECM signature in *KMT5B*-deficient DIPG cells compared to wild-type F8. The curves show the enrichment score on the y axis and the rank list metric on the x axis. Alongside is a heatmap representation of expression of significantly differentially expressed genes in the signature in all KMT5B/C-deficient DIPG cells compared to wild-type F8. (F) GSEA enrichment plots for an integrin signature in *KMT5B*-deficient DIPG cells compared to wild-type F8. The curves show the enrichment score on the y axis and the rank list metric on the x axis. Alongside is a heatmap representation of expression of significantly differentially expressed genes in the signature in all KMT5B/C-deficient DIPG cells compared to wild-type F8. (G) Immunofluorescence of DIPG cells invading through matrigel, stained for H3K20me1, K4K20me2 or H4K20me3, with DAPI as a counterstain. Scale bar is 200μm. (H) Light microscopy image of *KMT5B* wild-type, H3.3 K27M F8 cells and *KMT5B*-deficient F10 and K5B cells invading through matrigel. over 6 days. Scale bar is 1mm. (I) Quantitation of invasion area (y axis) over time (x axis) for *KMT5B* wild-type, H3.3 K27M F8 cells (grey) and *KMT5B*-deficient F10 and K5B cells (blue). Stats - ***p<0.001. (J) Overview of secretome experiment, whereby conditioned media was obtained from *KMT5B* wild-type, H3.3 K27M F8 cells (grey) and *KMT5B*-deficient F10 and K5B cells (blue) after 24 hours after removal of growth factors, and analysed by LC-MS/MS. A scaled Venn diagram shows the overlap in secreted proteins between the cells. (K) Heatmap of protein abundance for those identified as differentially secreted by F10 and K5B cells compared to F8, coloured by key provided. (L) Gene ontology enrichment analysis for proteins differentially secreted by *KMT5B*-deficient F10 cells compared to wild-type F8, plotted in bars according to their LOD score.

We next sought to explore whether the increased invasive capacity was a result of secreted factors produced in the KMT5B-deficient cells, and performed a comprehensive secretome analysis by LC-MS/MS (Figure 4J), identifying 170 differentially secreted proteins in F10 and/or K5B (Supplementary Table S4). Although there were common proteins identified in the two KMT5B-deficient models, the majority of these were unique to F10, which has diverged *in vitro*, and plays a role in signalling to F8 cells in heterogeneous cultures (Vinci et al., 2018). Proteins observed in both the lines included know secreted factors previously associated with a more invasive phenotype in this context, such as MMP2, VCAN, SERPINH1, ITGAV and IGFBP3 (Figure 4K), and were strongly enriched in processes linked to extracellular matrix re-organisation/degradation and cell adhesion, amongst others (Figure 4L).

### Changes in chromatin accessibility drives gene expression via distinct cis-regulatory elements

Having determined the profound and functionally relevant effects caused by *KMT5B/C* loss in DIPG through bulk profiling, we next explored the differences in our model systems at the single cell level. Using UMAP projections from 10X expression data (n = 16552 cells, median = 2798 per library), we were able to discretely resolve the libraries not only from B193 and the single cell-derived F10 culture, but also our KMT5B/C KD cells from the parental F8 cells, particularly the double-KD (Figure 5A). Overlaying gene expression signatures relating to H3K27me3 targets (Figure 5B), we observed a significant over-expression in *KMT5B/C*-deficient cells compared to F8, in keeping with bulk profiling (Figure 5C), though not homogenous across all cells from our models. This heterogeneity was particularly pronounced in relation to gene signatures associated with glioma cell identity (Neftel et al., 2019). As well as the substantial plasticity between mesenchymal (MES), astrocytic (AC), oligodendrocyte precursor-like (OPC) and neural precursor-like (NPC) cell states observed in our unselected B193 glioma stem-like cell culture, we also observed marked differences in distinct cell subpopulations in our KMT5B and/or KMT5C-KD clones (Supplementary Figure S5A). In particular, there was a shift in some cells towards OPC and/or NPC-like identity associated with the loss of the H4 methyltransferases (p<0.0001, ANOVA) (Supplementary Figure S5B). Of note, inferred DNA copy number profiles derived from 10X Genomics single-cell RNA sequencing (scRNA-seq), demonstrated largely consistent genomic profiles of our CRISPR-KD models, though some divergence such as an absence of 1q gain in F10 (Supplementary Figure S5C).

**Figure 5.**
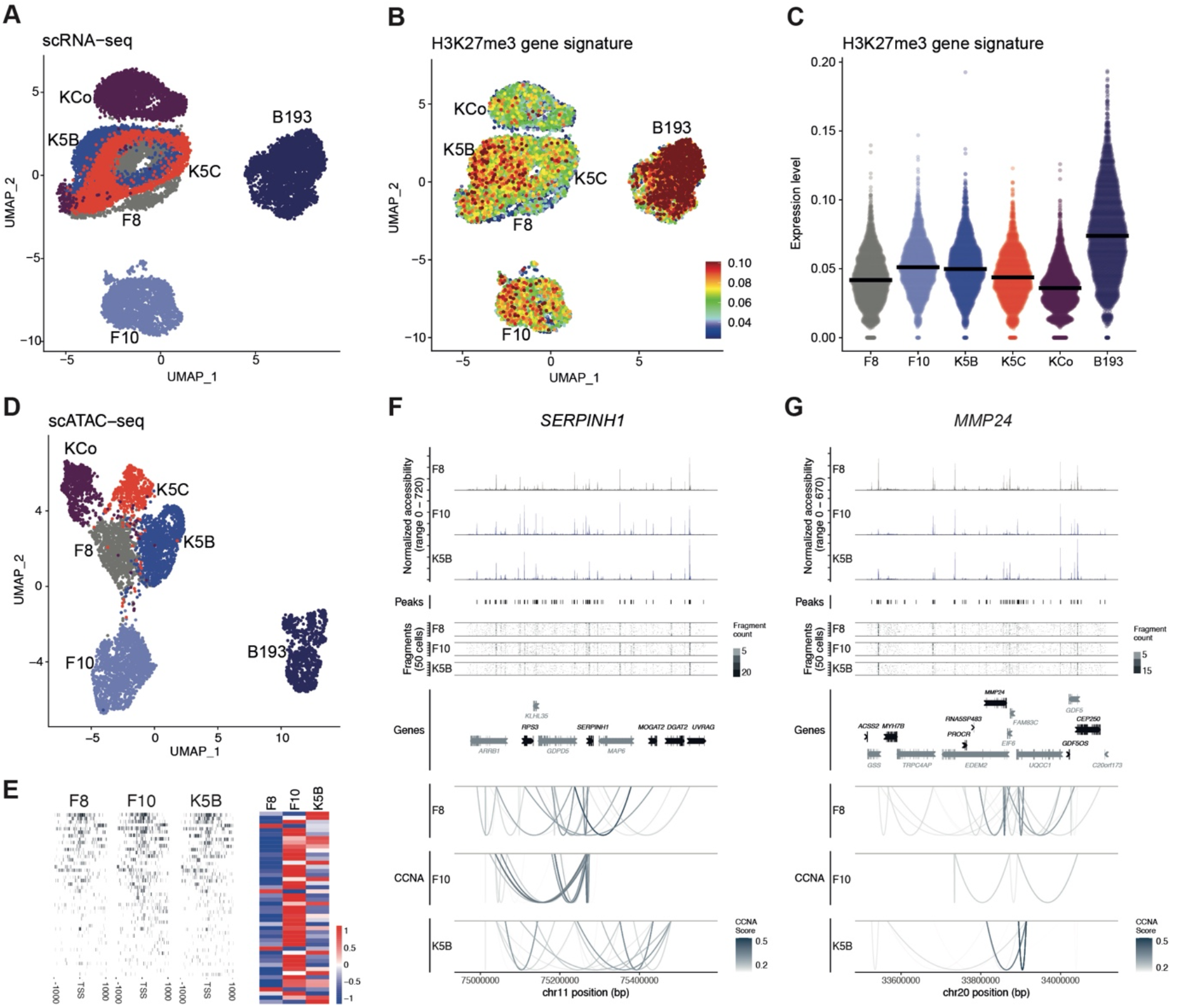
Chromatin accessibility and gene expression changes at the single cell level in response to KMT5B/C loss. (A) UMAP projection of 16552 single cells profiled for gene expression by RNA sequencing using the 10X platform, using the most variable genes across the libraries, and coloured by cell library. *KMT5B/C* wild-type F8, grey; *KMT5B*_R187* F10, light blue; *KMT5B*-KD K5B, blue; *KMT5C*-KD, red; *KMT5B/C* double-KD, purple; *KMT5B*_M646fs B193, dark blue. (B) Overlay of gene signature expression levels for H3K27me3 targets on the UMAP projection in (A), coloured according to the key provided. (C) Violin plots of the median gene expression of the H3K27me3 target signatures shown in (E), separated by library. The horizontal line represents the median across single cells, ****p<0.0001, ANOVA. (D) UMAP projection of 6310 single cells profiled by ATAC-seq using the 10X platform, using the most variable accessibilities across the libraries, and coloured by cell library. *KMT5B/C* wild-type F8, grey; *KMT5B*_R187* F10, light blue; *KMT5B*-KD K5B, blue; *KMT5C*-KD, red; *KMT5B/C* double-KD, purple; *KMT5B*_M646fs B193, dark blue. Fragment coverage of regions 1000bp up/downstream of the TSS (left) normalised to total reads per cell and library size (right) for the H3K27me3 signature from scATAC-seq data. (F) Cicero co-accessibility map for the region 300 kbp up- and downstream of *SERPINH1*, with all protein-coding genes in the region and accessibility peaks defined by Cellranger annotated. Co-accessibility is shown for each library with minimum threshold of 0.2. Per-library down-sampled accessibility and fragment coverage for 50 cells by 1500 bp bins are provided. (G) Cicero co-accessibility map for the region 300 kbp up- and downstream of *MMP24*, with all protein-coding genes in the region and accessibility peaks defined by Cellranger annotated. Co-accessibility is shown for each library with minimum threshold of 0.2. Per-library down-sampled accessibility and fragment coverage for 50 cells by 1500 bp bins are provided.

To determine the extent to which these changes were associated with chromatin dynamics, we carried out single cell ATAC-seq, also using the 10X platform (n = 6310 cells, median = 1020 per library). Here the libraries were resolved to an even greater extent by UMAP projection, particularly among the single gene KD cells (Figure 5D). Again focusing on H3K27me3 targets, we observed a distinct increase in open chromatin in F10 and K5B compared to F8 (Figure 5E), which could also be observed more globally (Supplementary Figure S5D). Most strikingly we observed key differences in co-accessibility of *cis*-regulatory domains associated with *KMT5B* loss. Using Cicero (Pliner et al., 2018) to identify co-accessible pairs of DNA elements, we identified numerous differentially expressed genes in F10 and K5B compared to F8 to be associated with open chromatin at distinct regulatory elements which seemingly account for their transcriptional activation. These included key genes from our previously identified gene signatures associated with mesenchymal transition, such as *SERPINH1* (Figure 5F) and *MMP24* (Figure 5G). These and similarly differentially *cis-* regulated genes demonstrated heterogenous accessibility profiles at the single cell level, demonstrating the wide-ranging effects on chromatin architecture driving differential expression wrought by KMT5B loss (Supplementary Table S5).

### KMT5B loss drives transcriptional diversity, and confers poor prognosis in paediatric high grade glioma

Finally, we further carried out SmartSeq2 single cell RNA sequencing in order to assess transcriptional heterogeneity by examining full-length transcripts. These data, although derived from fewer cells than 10X (n=746, median=131 per library), were able to discriminate the various cell model libraries to a similar extent (Supplementary Figure S6A), and showed similar patterns of inferred DNA copy number change (Supplementary Figure S6B). Applying the GINI index to the expression data reinforced the transcriptional heterogeneity in this sample compared to the clones, but also demonstrated a shift towards a more diverse transcriptional profile in *KMT5B/C*-deficient cells compared to F8 (Figure 6A). This was even more apparent by calculating Shannon diversity indices for the different libraries, with the specifically *KMT5B*-deficient libraries (and not *KMT5C*) showing significantly more heterogeneity than the wild-type (p< 0.001 for F10, K5B, KCo, B193 compared to F8, t-test) (Figure 6B). This was found to be particularly true for certain key genes, such as *NCAN*, and *VGF*, in which expression was markedly different across cellular subpopulations, unlike housekeeping genes such as *GAPDH* provided as a comparator (Figure 6C) (Supplementary Table S6).

**Figure 6.**
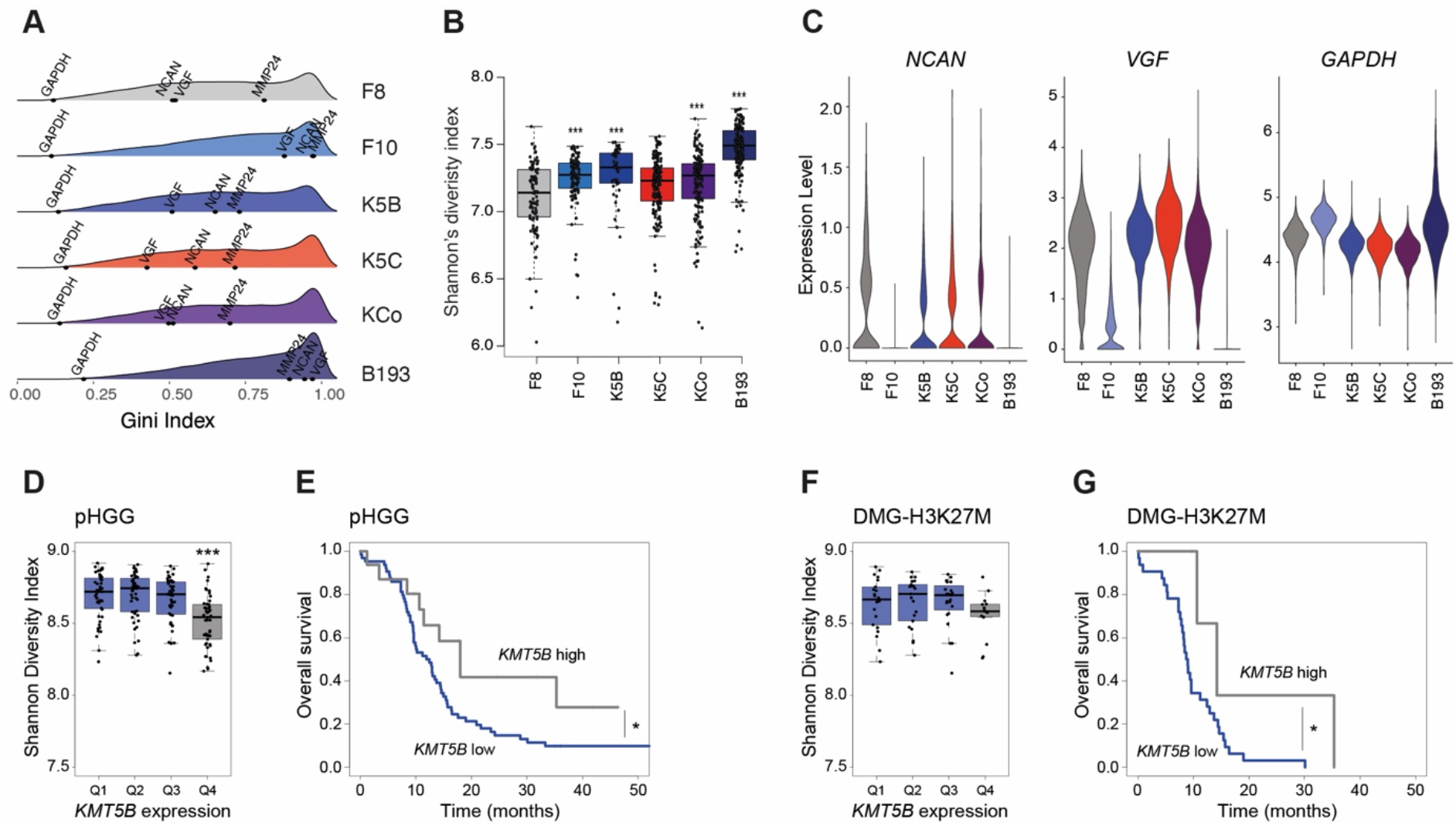
KMT5B loss drives transcriptional heterogeneity and confers shorter survival in paediatric high grade glioma. (A) Density plot of GINI indices calculated from SmartSeq2 single-cell RNA-seq data, separated by cell library, and with selected genes labelled. *KMT5B/C* wild-type F8, grey; *KMT5B*_R187* F10, light blue; *KMT5B*-KD K5B, blue; *KMT5C*-KD, red; *KMT5B/C* double-KD, purple; *KMT5B*_M646fs B193, dark blue. (B) Boxplot of Shannon’s diversity indices calculated from SmartSeq2 single-cell RNA-seq data, separated by cell library, The horizontal line with the boxes represents the median, the box the interquartile range, and the whiskers spanning 95% of the data. *** p<0.0001, t-test. *KMT5B/C* wild-type F8, grey; *KMT5B*_R187* F10, light blue; *KMT5B*-KD K5B, blue; *KMT5C*-KD, red; *KMT5B/C* double-KD, purple; *KMT5B*_M646fs B193, dark blue. (C) Violin plot of gene expression from SmartSeq2 single-cell RNA-seq data, separated by cell library, for selected genes based upon transcriptional diversity differences. (D) Boxplot of Shannon’s diversity indices calculated bulk RNA-seq data from published paediatric high grade glioma tumours, separated by *KMT5B* expression. The horizontal line with the boxes represents the median, the box the interquartile range, and the whiskers spanning 95% of the data. *** p<0.0001, t-test. *KMT5B* low, blue; *KMT5B* high, grey. (E) Survival curve for paediatric high grade glioma, separated by *KMT5B* expression levels. Time in months is given on the x axis, overall survival proportions are provided on the y axis. * p<0.05, log-rank test. (F) Boxplot of Shannon’s diversity indices calculated bulk RNA-seq data from published H3K27M-DMG tumours, separated by *KMT5B* expression. The horizontal line with the boxes represents the median, the box the interquartile range, and the whiskers spanning 95% of the data. *** p<0.0001, t-test. *KMT5B* low, blue; *KMT5B* high, grey. (G) Survival curve for H3K27M-DMG, separated by *KMT5B* expression levels. Time in months is given on the x axis, overall survival proportions are provided on the y axis. * p<0.05, log-rank test.

To determine whether this association held true in patient tumour samples, we reverted to publicly available bulk RNAseq data, and explored the associations between *KMT5B* expression and Shannon diversity. Across all paediatric high grade glioma, there was a significant reduction in transcriptional heterogeneity in samples with high levels of *KMT5B* (p>0.0001, t-test) (Figure 6D), recapitulating the single-cell data. Importantly, paediatric high grade glioma with lower levels of *KMT5B* had a significantly poorer prognosis than those with high expression (p=0.0126, log-rank test) (Figure 6E). Although the numbers were smaller, the same held true when considering H3K27M-DMG alone (p=0.05, log-rank test) (Figure 6F,G), demonstrating the importance of transcriptional heterogeneity in providing the evolutionary substrate for overcoming therapeutic interventions. This was again in contrast to *KMT5C*, whereby diversity increased with higher expression, and there was no significant impact on clinical outcome (Supplementary Figure S6C-F).

In summary, we have identified a novel mechanism whereby loss of an H4 methyltransferase results in changes in chromatin accessibility which conveys an ablation of H3K27me3 binding at bivalent loci otherwise retained in H3K27M DIPG cells. This results in profound transcriptional changes associated with pro-tumorigenic processes including mesenchymal transition, invasion and cellular heterogeneity, and highlights the role of transhistone mechanisms in these childhood brain tumours of unmet clinical need (Figure 7).

**Figure 7.**
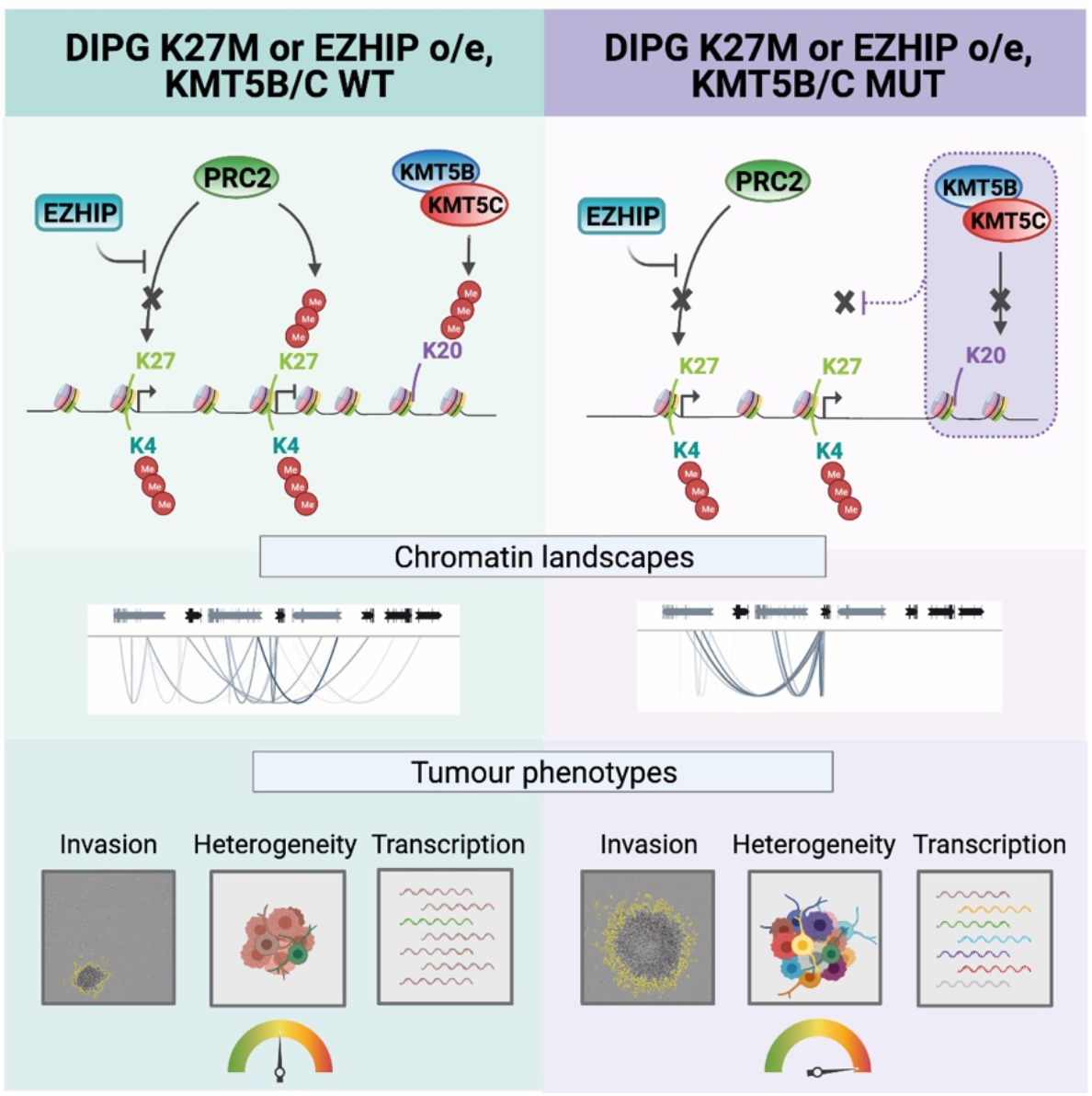
Proposed mechanism of action. *KMT5B/C* loss leads to focal loss of H4K20me2/3 in addition to loss of H3K27me3 at loci otherwise retained in H3K27M/EZHIP tumours (top panel). This has profound consequential changes on chromatin accessibility and gene expression (middle panel), leading to enhanced transcriptional heterogeneity, cell plasticity, and pro-tumorigenic features such as invasiveness (bottom panel).

## DISCUSSION

H3K27M DMG/DIPG maintain a cellular hierarchy reflecting the stalled developmental origins of these tumour subtypes (Filbin et al., 2018; Jessa et al., 2019). We have shown the genotypic and phenotypic divergence of these cancer stem-like cells, whereby subclones derived from them have distinct tumorigenic capabilities acting in concert; the heterogeneity itself appears to be the evolutionary unit of selection (Little et al., 2012; Vinci et al., 2018). Specific subclones acquire unique genetic aberrations, and generate and respond to diverse signals into the tumour milieu - driving infiltration, early escape from the pons, and colonisation of distant metastatic sites (Qin et al., 2017; Venkatesh et al., 2015; Vinci et al., 2018).

Mechanistically, the manner in which these cells arise, and interact, is driven by a complex interplay between genetic and epigenetic mechanisms (Jones, 2020; Jones and Baker, 2014; Sturm et al., 2014). Genetic insults arising in precisely defined developmental contexts lock in profound changes in chromosomal architecture, chromatin accessibility and gene expression, which defines cellular states (Filbin et al., 2018; Nagaraja et al., 2017). Here for the first time, we begin to unravel the mechanism by which this takes place, whereby loss-of-function mutations in histone H4 modifying enzymes cause profound changes in gene expression via altered chromatin accessibility and H3K27me3 binding.

The lysine methyltransferases KMT5B (SUV420H1) and KMT5C (SUV420H2) catalyse H4K20 di- and tri-methylation, a post-translational modification associated with constitutive heterochromatin, telomeres and centromeres (Brohm et al., 2019); it is also involved with euchromatic gene silencing, and plays a key role in the maintenance of genomic integrity via DNA repair, replication and chromatin compaction (Jorgensen et al., 2013; Schotta et al., 2008). Although originally described in subclonal proportions (Schwartzentruber et al., 2012; Vinci et al., 2018), our identification of a DIPG sample with a clonal truncating mutation in *KMT5B* demonstrates the importance of this novel tumour suppressor gene in this context.

In our ‘natural’ isogenic subclones, engineered KD cells, and in bulk clonally mutant cultures, we observe the rewiring of the epigenome caused by loss of the H4 methyltransferase in the background of H3K27M / EZHIP-overexpressing DIPG cells. A key feature of these tumours is the global loss of the repressive H3K27me3 mark, though the retention of this modification at key loci has been shown to regulate key oncogenic pathways (Bender et al., 2013; Chan et al., 2013; Funato et al., 2014; Larson et al., 2019; Mohammad et al., 2017). These retained loci were found to be consistent across individuals tumour specimens, with our data suggestive of an H4-related mechanism maintaining the modifications. Loss of this additional fine-control appears to drive profound transcriptional changes promoting tumorigenesis, particularly in relation to cell identify, mesenchymal transition, and invasion/migration; even if only occurring in a subset of tumour cells, a co-operative advantage may be conferred on neighbouring cells through the upregulation and exocytosis of important secreted factors (Vinci et al., 2018).

The importance of such trans-histone effects in cancer is only recently garnering attention, some 20 years after the discovery of a ‘trans-tail’ regulation of histone modification, linking H2B ubiquitination to H3 methylation and gene silencing in yeast (Briggs et al., 2002; Sun and Allis, 2002). Whilst most attention to this cross-talk has focussed on the links between H2B ubiquitination and H3K79 trimethylation by DOT1L (Anderson et al., 2019; Valencia-Sanchez et al., 2021; Valencia-Sanchez et al., 2019; Worden et al., 2019), H3K4me3 has also been implicated via COMPASS (Hsu et al., 2019). The H3K4 methyltransferases MLL3/4 have been shown to bind to histone H4 during stem cell differentiation (Liu et al., 2019), and moreover MLL1-mediated histone H3K4 methylation has been found to act synergistically with histone H4K16 via the histone acetyltransferase MOF (Dou et al., 2005).

H4K20me3 has been found to co-localise with H3K9me3 in proliferating myoblasts (Bhanu et al., 2019) and ES cells (Xu and Kidder, 2018), whilst bivalent chromatin domains consisting of H3K4me3 and H3K27me3, enriched at developmental genes that are repressed in ES cells but active during differentiation, were found to have a simultaneous presence of H4K20me3 (Liu et al., 2019). A comparative genomics approach highlighted a substantial set of genes enriched with H3K27me3 and H4K20me3 in baboon subventricular zone (SVZ)-derived NSCs are altered in human GBM, suggesting a mechanism to control gene expression and proliferation in the progenitor cell compartment, protecting against abnormal cell cycle entry (Rhodes et al., 2016).

The profound shift in co-accessibility of DNA elements observed with *KMT5B/C* loss demonstrates the complexity of *cis*-regulation in these tumours. Heterogenous chromatin landscapes have recently been shown to play an important role in glioma stem-like cell diversity in adult glioblastoma (Guilhamon et al., 2021), whilst in H3G34R/V mutant glioma, it was recently shown that the specific lineage context of these tumours may facilitate PDGFRA co-option through a chromatin loop connecting *PDGFRA* to *GSX2* regulatory elements, promoting *PDGFRA* overexpression (Chen et al., 2020). A fundamental consequence of the altered gene expression landscape observed with *KMT5B* loss is a significant increase in the transcriptional heterogeneity, in patient samples as well as model systems, as has also been observed with the histone demethylase *KDM5B* in breast cancer (Hinohara et al., 2018). This likely provides the substrate for cellular phenotypic diversity and treatment escape, and confers the poor prognosis found in patients with these alterations.

These data add a layer of complexity to devising treatment strategies for these oncohistone-driven tumours, but also new opportunities. In addition to identifying distinct therapeutic vulnerabilities in subclones with *KMT5B* loss, or devising strategies to disrupt communication between different tumour cell subpopulations, epigenetic modifying agents used in combination with targeted inhibitors or other approaches may help counteract the transcriptional changes driving the poor clinical response characteristic of these children’s tumours.

## ACKNOWLEDGEMENTS

This work was supported by Cancer Research UK (C13468/A23536), Children with Cancer UK (16-212 and 16-234), Billie Butterfly Fund and the Brain Tumour Charity, the CRIS Cancer Foundation and the DIPG Collaborative. A.S. is supported by the Wellcome Trust (202778/B/16/Z) and Cancer Research UK (A22909). We acknowledge funding from the National Institute of Health (NCI U54 CA217376) to A.S. This work was also supported a Wellcome Trust award to the Centre for Evolution and Cancer (105104/Z/14/Z).

## AUTHOR CONTRIBUTIONS

KK, MV and CJ conceived the study. KK, VM, AB, ST, EY-R, IL and LY carried out experiments. DC provided data. AM and YG performed bioinformatic analyses. KK, AM, YB, HT, JC, AS, MF and CJ analysed data. KK and CJ wrote the manuscript. All authors read and approved the manuscript.

## DECLARATION OF INTERESTS

None

## SUPPLEMENTAL FIGURES

**Supplementary Figure S1.**
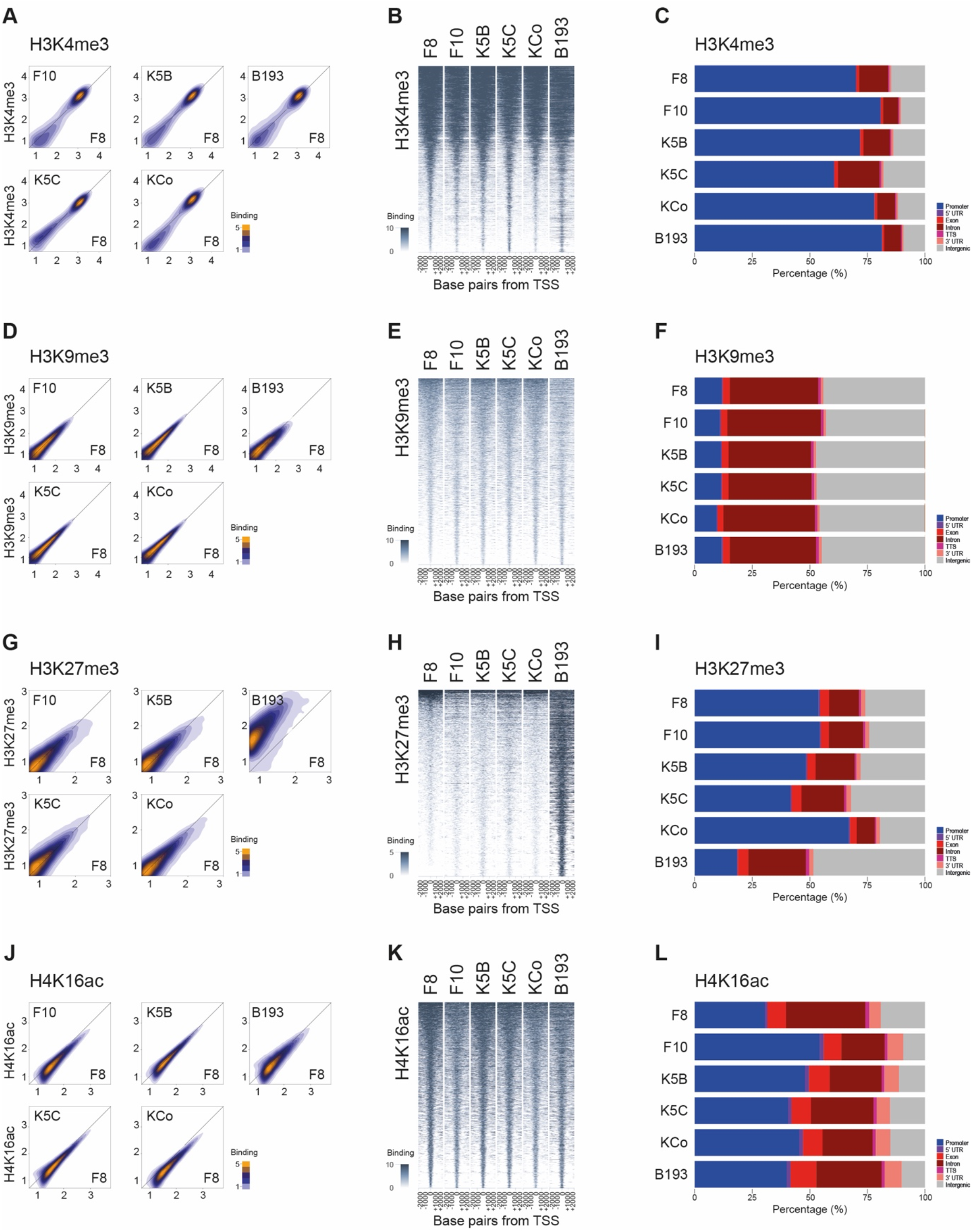
Binding of key histone H3/H4 marks in KMT5B/C-deficient patient-derived DIPG cells. (A) Density plot of binding of H3K4me3 peaks as assessed by ChIP-seq in *KMT5B/C*-deficient cells (y axes) compared to wild-type F8 cells (x axes). Density of peaks is coloured according to the key provided. (B) Heatmap of H3K4me3 binding in DIPG cells ranked ordered by reads, and plotted spanning 2kb either side of the transcriptional start site. Extent of binding is coloured according to the key provided. (C) Stacked barplot showing annotated distribution of H3K4me3 binding in all cells in distinct genic and intergenic regions, coloured according to the key provided and plotted as an overall percentage per cell. (D) Density plot of binding of H3K9me3 peaks as assessed by ChIP-seq in *KMT5B/C*-deficient cells (y axes) compared to wild-type F8 cells (x axes). Density of peaks is coloured according to the key provided. (E) Heatmap of H3K9me3 binding in DIPG cells ranked ordered by reads, and plotted spanning 2kb either side of the transcriptional start site. Extent of binding is coloured according to the key provided. (F) Stacked barplot showing annotated distribution of H3K9me3 binding in all cells in distinct genic and intergenic regions, coloured according to the key provided and plotted as an overall percentage per cell. (G) Density plot of binding of H3K27me3 peaks as assessed by ChIP-seq in *KMT5B/C*-deficient cells (y axes) compared to wild-type F8 cells (x axes). Density of peaks is coloured according to the key provided. (H) Heatmap of H3K27me3 binding in DIPG cells ranked ordered by reads, and plotted spanning 2kb either side of the transcriptional start site. Extent of binding is coloured according to the key provided. (I) Stacked barplot showing annotated distribution of H3K27me3 binding in all cells in distinct genic and intergenic regions, coloured according to the key provided and plotted as an overall percentage per cell. (J) Density plot of binding of H4K16ac peaks as assessed by ChIP-seq in *KMT5B/C*-deficient cells (y axes) compared to wild-type F8 cells (x axes). Density of peaks is coloured according to the key provided. (K) Heatmap of H4K16ac binding in DIPG cells ranked ordered by reads, and plotted spanning 2kb either side of the transcriptional start site. Extent of binding is coloured according to the key provided. (K) Stacked barplot showing annotated distribution of H4K16ac binding in all cells in distinct genic and intergenic regions, coloured according to the key provided and plotted as an overall percentage per cell.

**Supplementary Figure S2.**
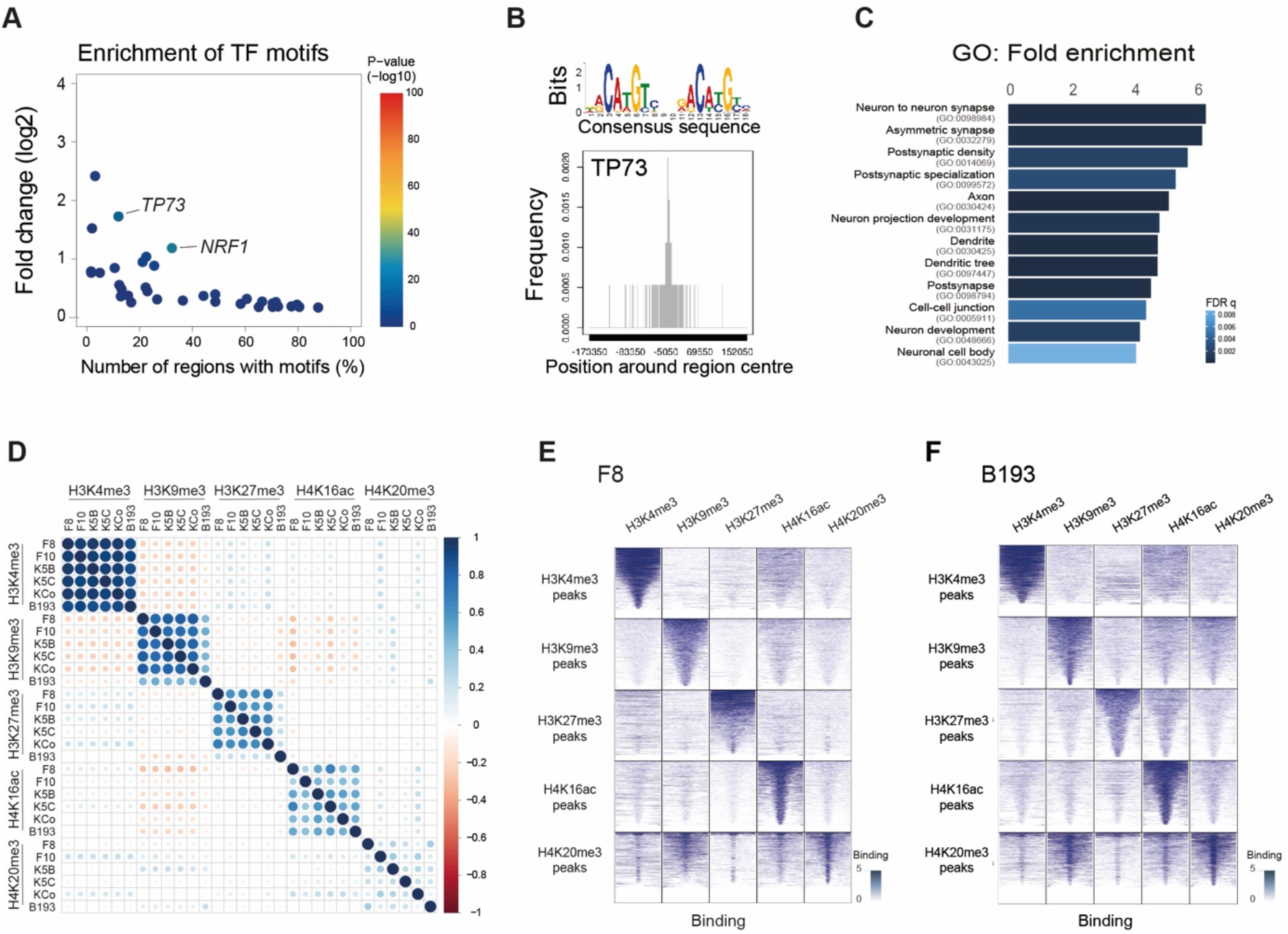
H4K20me3 binding and correlations of H3/H4 modifications in KMT5B/C-deficient DIPG cells. (A) Transcription factor (TF) enrichment for H4K20me3 bound loci in *KMT5B/C* wild-type F8 cells, plotted as percentage of motif-containing regions (x axis) against log2 fold change compared to control genomic regions (y axis), and coloured by −log10 p value (t-test) according to the scale provided. TF motifs with p value < 1×10^−10^ are labelled. Consensus binding sequence of the enriched *TP73* motif, with its location in base pairs around the centre of the H4K20me3 bound region (x axis) plotted against frequency (y axis). Gene ontology enrichment analysis for genes bound by H4K20me3 in *KMT5B/C* wild-type F8 cells, plotted in bars according to their fold enrichment, coloured by q value calculated as the false discovery rate (FDR q) according to the scale provided. (D) Global correlation plot of all H3K4me3, H3K9me3, H3K27me3, H4K16ac and H4K20me3 marks across all cells. Size and colour of circle is dependent on the Pearson’s correlation coefficient, according to the scale provided. (E) Heatmap of binding in *KMT5B* wild-type, H3.3 K27M F8 cells for H3K4me3, H3K9me3, H3K27me3, H4K16ac and H4K20me3, ranked ordered by reads, and plotted spanning 2kb either side of the transcriptional start site. Extent of binding is coloured according to the key provided. (F) Heatmap of binding in *KMT5B*_M646fs, *EZHIP*-overexpressing B193 cells for H3K4me3, H3K9me3, H3K27me3, H4K16ac and H4K20me3, ranked ordered by reads, and plotted spanning 2kb either side of the transcriptional start site. Extent of binding is coloured according to the key provided.

**Supplementary Figure S3.**
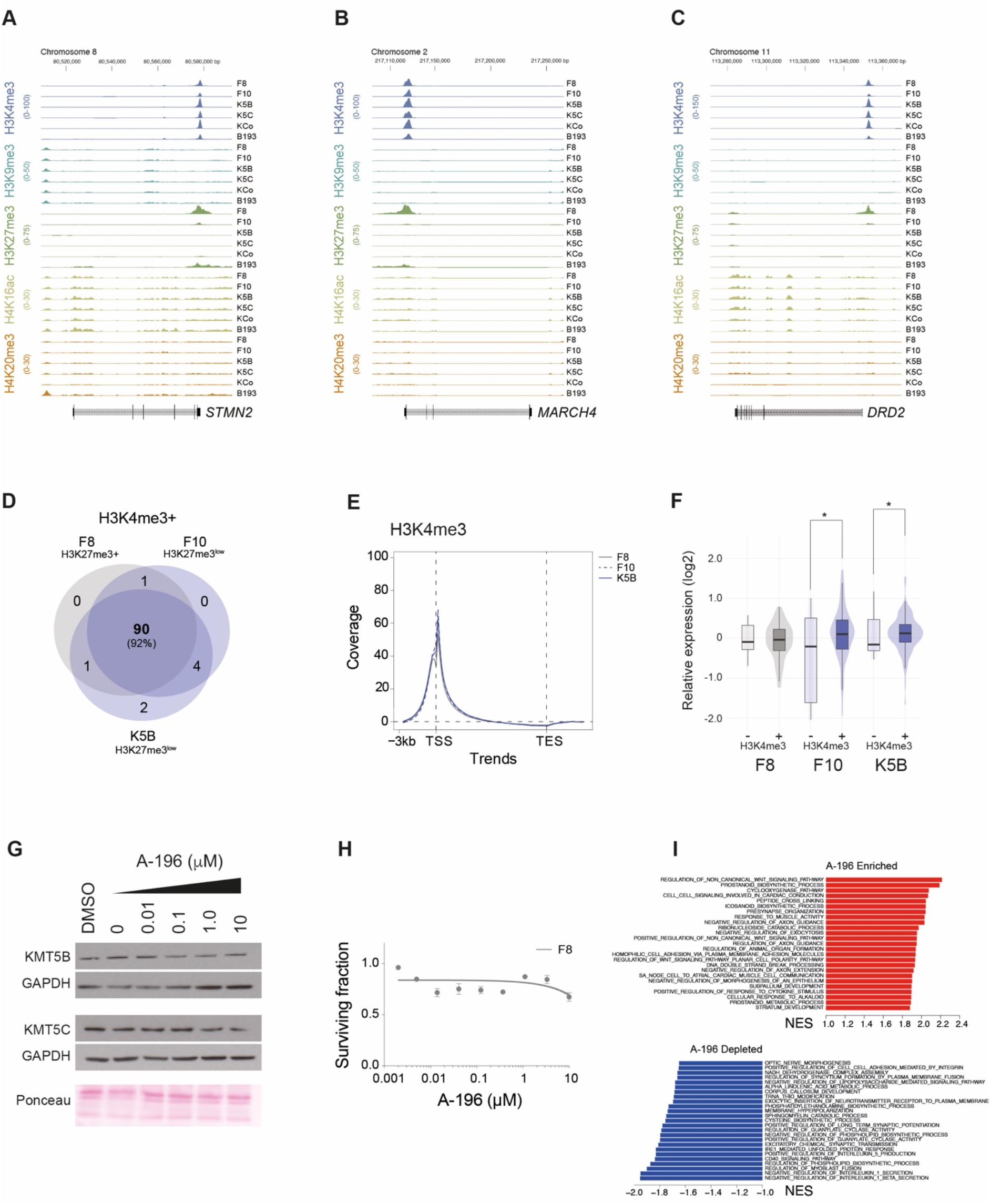
Bivalency and KMT5B-dependency of differently H3K27me3-bound and expressed genes in DIPG. (A) ChIP-seq coverage plot of H3K4me3 (blue), H3K9me3 (aqua), H3K27me3 (green), H4K16ac (mustard) and H4K20me3 (orange) occupancy plotted as read depth (y axes) for each DIPG cell model for the coding sequence of *STMN2* on chromosome 8, with coordinates given in base pairs (x axis) and the direction of transcription from the promoter indicated. Plots from all samples for any given mark are on the same scale. (B) ChIP-seq coverage plot of H3K4me3 (blue), H3K9me3 (aqua), H3K27me3 (green), H4K16ac (mustard) and H4K20me3 (orange) occupancy plotted as read depth (y axes) for each DIPG cell model for the coding sequence of *MARCH4* on chromosome 2, with coordinates given in base pairs (x axis) and the direction of transcription from the promoter indicated. Plots from all samples for any given mark are on the same scale. (C) ChIP-seq coverage plot of H3K4me3 (blue), H3K9me3 (aqua), H3K27me3 (green), H4K16ac (mustard) and H4K20me3 (orange) occupancy plotted as read depth (y axes) for each DIPG cell model for the coding sequence of *DRD2* on chromosome 11, with coordinates given in base pairs (x axis) and the direction of transcription from the promoter indicated. Plots from all samples for any given mark are on the same scale. (D) Venn diagram of H3K4me3 bound loci which are concurrently bound by H3K27me3 in *KMT5B* wild-type, H3.3 K27M F8 cells (grey) and lost in *KMT5B*-deficient F10 and K5B cells (blue). (E) Trend plots of coverage for H3K4me3 binding relative to +/− 3kb either side of the transcriptional start sites (TSS) and transcriptional end sites (TES) of all genes for *KMT5B* wild-type F8 and -deficient F10 and K5B cells. (F) Violin plot of mRNA expression for genes within the highly H3K27me3-bound loci in *KMT5B* wild-type, H3.3 K27M F8 cells, for F8 (grey) and *KMT5B*-deficient F10 and K5B cells (blue), split by loci with and without H3K4me3 binding. The horizontal line with the boxes represents the median, the box the interquartile range, and the whiskers spanning 95% of the data. * p<0.05, t-test. (G) Western blot for KMT5B and KMT5C, with individual GAPDH and/or Ponceau staining as loading controls, for *KMT5B* wild-type, H3.3 K27M F8 cells in response to increasing concentrations of the dual KMT5B/C inhibitor A-196. (H) Cell viability of *KMT5B* wild-type, H3.3 K27M F8 cells, plotted as surviving fraction (y axis) in response to increasing concentrations of the dual KMT5B/C inhibitor A-196 (x axis). (I) GSEA analysis differential expression by RNA-seq of F8 cells treated with 1μM A-196 for 24 hours. Normalized enrichment scores (NES) are plotted for those gene sets enriched (top, red) or depleted (bottom, blue) upon drug treatment.

**Supplementary Figure S4.**
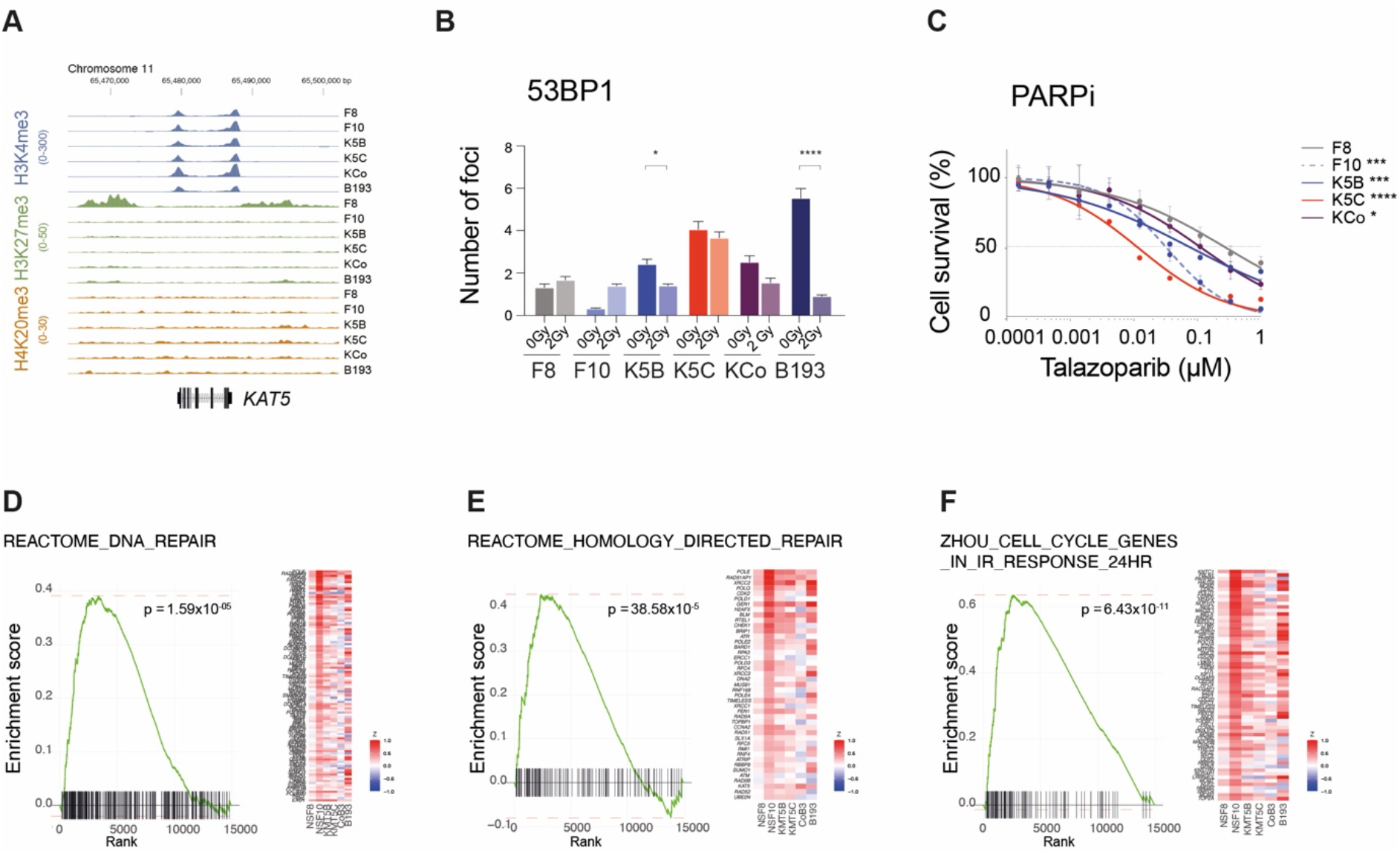
Loss of KMT5B causes gene expression changes promoting DNA repair abnormalities via H3K27me3/H3K4me3. (A) ChIP-seq coverage plot of H3K4me3 (blue), H3K27me3 (green) and H4K20me3 (orange) occupancy plotted as read depth (y axes) for each DIPG cell model for the coding sequence of *KAT5* on chromosome 1, with coordinates given in base pairs (x axis) and the direction of transcription from the promoter indicated. Plots from all samples for any given mark are on the same scale. (B) Barplot of quantitation of 53BP1 foci for DIPG cells after 2 Gy radiation. *KMT5B/C* wild-type F8, grey; *KMT5B*_R187* F10, light blue; *KMT5B*-KD K5B, blue; *KMT5C*-KD, red; *KMT5B/C* double-KD, purple; *KMT5B*_M646fs B193, dark blue. ****p<0.0001, *p<0.05, ANOVA. (C) Cell viability of DIPG cells plotted as surviving fraction (y axis) in response to increasing concentrations of the PARP inhibitor talazoparib (x axis). *KMT5B/C* wild-type F8, grey; *KMT5B*_R187* F10, light blue; *KMT5B*-KD K5B, blue; *KMT5C*-KD, red; *KMT5B/C* double-KD, purple. ****p<0.0001, ***p<0.001, *p<0.05, ANOVA. (D) GSEA enrichment plots for a DNA repair signature in *KMT5B*-deficient DIPG cells compared to wild-type F8. The curves show the enrichment score on the y axis and the rank list metric on the x axis. Alongside is a heatmap representation of expression of significantly differentially expressed genes in the signature in all KMT5B/C-deficient DIPG cells compared to wild-type F8. (E) GSEA enrichment plots for a homology-directed repair signature in *KMT5B*-deficient DIPG cells compared to wild-type F8. The curves show the enrichment score on the y axis and the rank list metric on the x axis. Alongside is a heatmap representation of expression of significantly differentially expressed genes in the signature in all KMT5B/C-deficient DIPG cells compared to wild-type F8. (F) GSEA enrichment plots for an IR-mediated cell cycle response signature in *KMT5B*-deficient DIPG cells compared to wild-type F8. The curves show the enrichment score on the y axis and the rank list metric on the x axis. Alongside is a heatmap representation of expression of significantly differentially expressed genes in the signature in all KMT5B/C-deficient DIPG cells compared to wild-type F8.

**Supplementary Figure S5.**
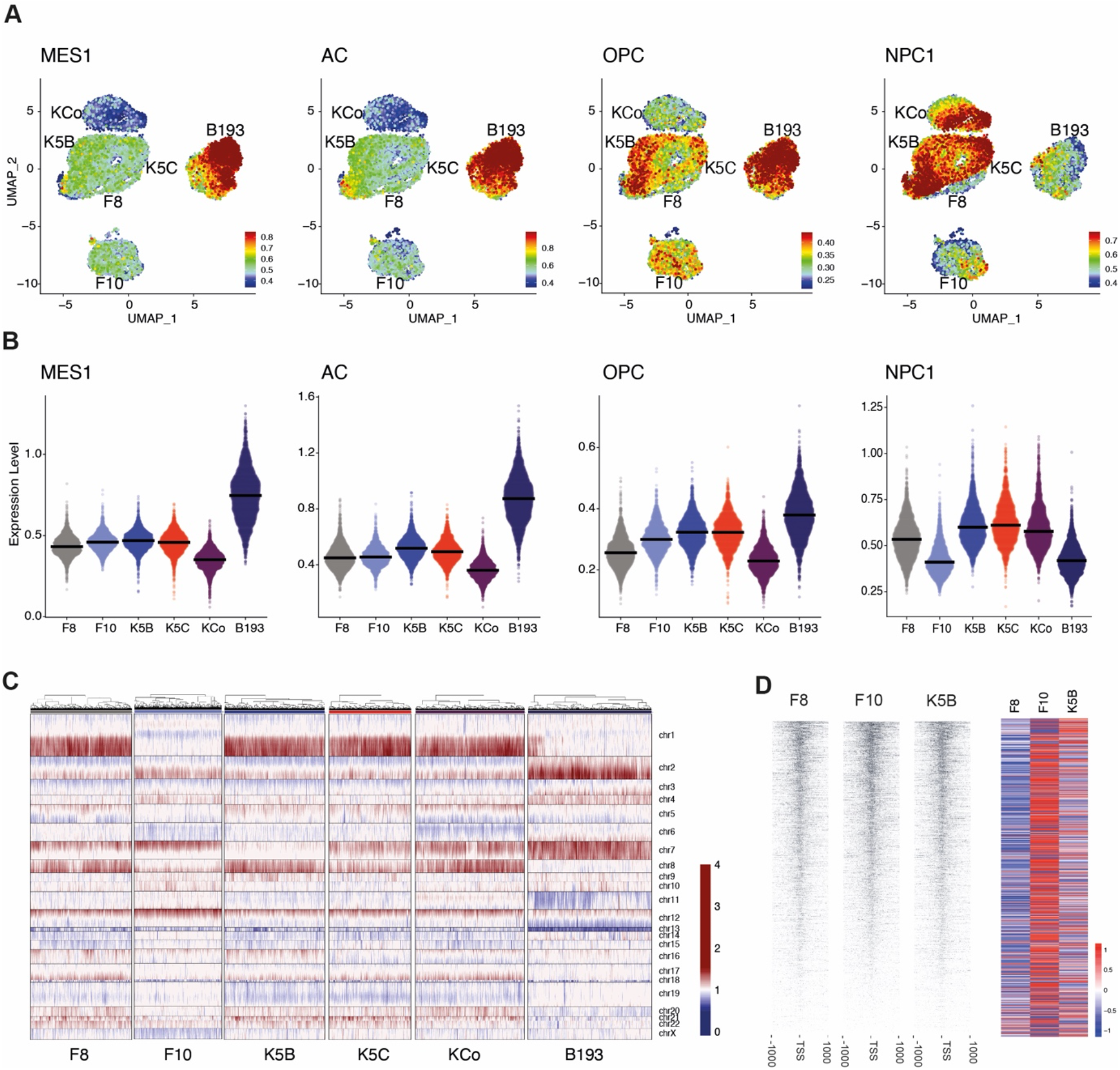
Single cell RNA-seq and ATAC-seq of DIPG cells and KMT5B/C-deficient clones. (A) UMAP projection of single cells profiled for gene expression by RNA sequencing using the 10X platform, coloured by the expression signatures representing mesenchymal-like cells (MES), astrocytes (AC), oligodendrocyte precursor cells (OPC) and neural progenitor cells (NPC), according to the scale provided. (B) Violin plots of the median gene expression of the MES, AC, OPC and NPC target signatures shown in (A), separated by library. *KMT5B/C* wild-type F8, grey; *KMT5B*_R187* F10, light blue; *KMT5B*-KD K5B, blue; *KMT5C*-KD, red; *KMT5B/C* double-KD, purple; *KMT5B*_M646fs B193, dark blue. The horizontal line represents the median across single cells, *** adjusted p value < 1×10^−15^; ** p < 1×10^−10^; * adjusted p value < 1×10^−5^. (C) Copy number heatmap derived from 10X single cell RNA-seq data, split by cell libraries. Columns represent individual cells, and rows genes arranged in genomic location from chromosome 1 to X. Red represents DNA copy number gain, blue represents loss, according to the scale provided. *KMT5B/C* wild-type F8, grey; *KMT5B*_R187* F10, light blue; *KMT5B*-KD K5B, blue; *KMT5C*-KD, red; *KMT5B/C* double-KD, purple; *KMT5B*_M646fs B193, dark blue. (D) Fragment coverage of regions 1000bp up/downstream of the TSS (left) normalised to total reads per cell and library size (right) for al genomic regions from scATAC-seq data.

**Supplementary Figure S6.**
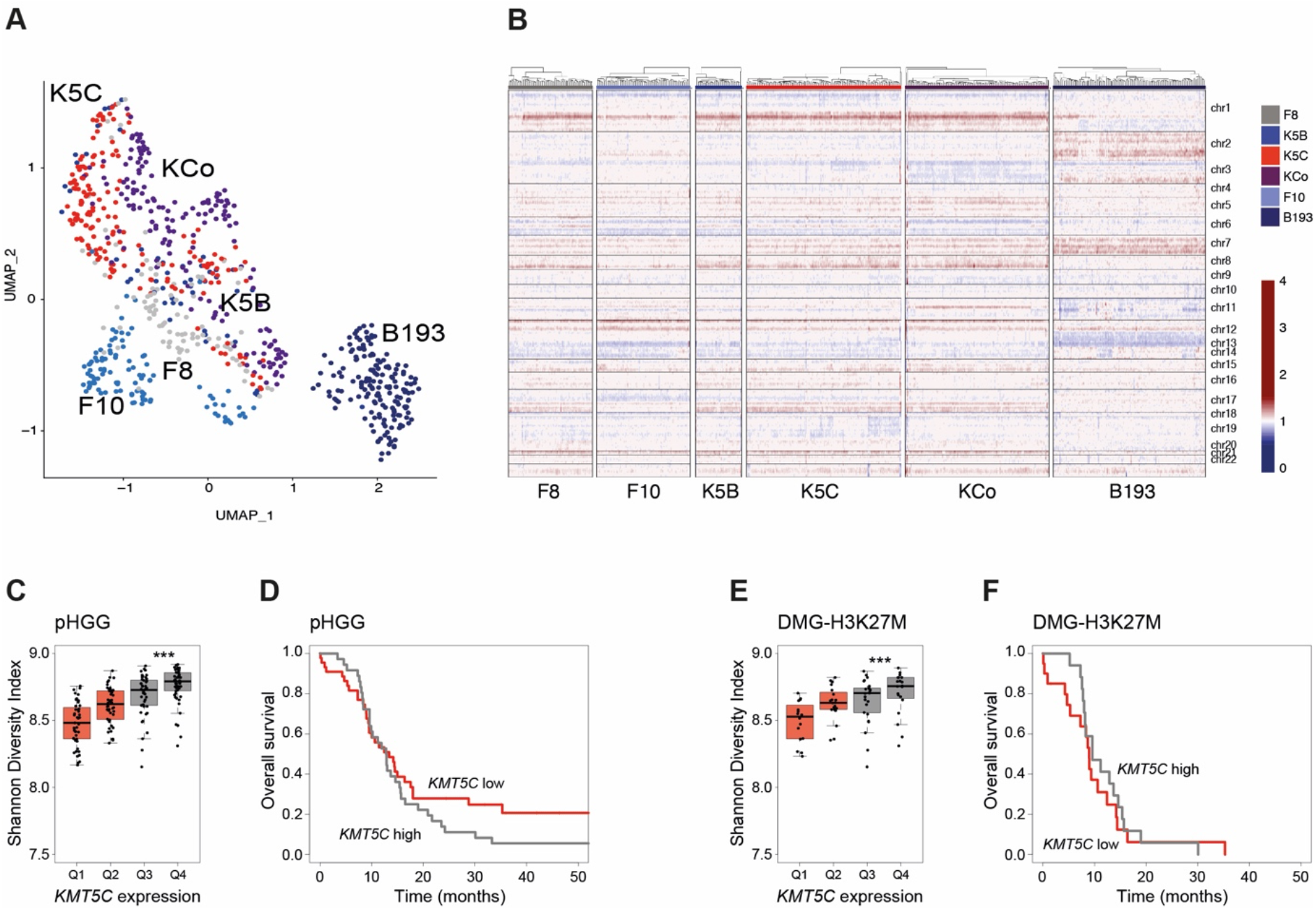
KMT5B loss drives transcriptional heterogeneity and confers shorter survival in paediatric high grade glioma. (A) UMAP projection of 746 single cells profiled for gene expression by RNA sequencing using the SmartSeq2 platform, using the most variable genes across the libraries, and coloured by cell library. *KMT5B/C* wild-type F8, grey; *KMT5B*_R187* F10, light blue; *KMT5B*-KD K5B, blue; *KMT5C*-KD, red; *KMT5B/C* double-KD, purple; *KMT5B*_M646fs B193, dark blue. (B) Copy number heatmap derived from SmartSeq2 single cell RNA-seq data, split by cell libraries. Columns represent individual cells, and rows genes arranged in genomic location from chromosome 1 to X. Red represents DNA copy number gain, blue represents loss, according to the scale provided. *KMT5B/C* wild-type F8, grey; *KMT5B*_R187* F10, light blue; *KMT5B*-KD K5B, blue; *KMT5C*-KD, red; *KMT5B/C* double-KD, purple; *KMT5B*_M646fs B193, dark blue. (C) Boxplot of Shannon’s diversity indices calculated bulk RNA-seq data from published paediatric high grade glioma tumours, separated by *KMT5C* expression. The horizontal line with the boxes represents the median, the box the interquartile range, and the whiskers spanning 95% of the data. *** p<0.0001, t-test. *KMT5C* low, red; *KMT5C* high, grey. (D) Survival curve for paediatric high grade glioma, separated by *KMT5C* expression levels. Time in months is given on the x axis, overall survival proportions are provided on the y axis. * p<0.05, log-rank test. (E) Boxplot of Shannon’s diversity indices calculated bulk RNA-seq data from published H3K27M-DMG tumours, separated by *KMT5C* expression. The horizontal line with the boxes represents the median, the box the interquartile range, and the whiskers spanning 95% of the data. *** p<0.0001, t-test. *KMT5C* low, red; *KMT5C* high, grey. (F) Survival curve for H3K27M-DMG, separated by *KMT5C* expression levels. Time in months is given on the x axis, overall survival proportions are provided on the y axis. * p<0.05, log-rank test.

## SUPPLEMENTAL TABLES

**Supplementary Table S1** – Differentially lost H4K20me3 peaks in *KMT5B/C*-deficient DIPG cells compared to wild-type, as assessed by ChIP-seq.

**Supplementary Table S2** – Shared retained H3K27me3 peaks between F8 cells and independent samples of H3.3K27M and H3.1K27M DIPG, as assessed by ChIP-seq.

**Supplementary Table S3** – H3K27me3 binding, as assessed by ChIP-seq, and gene expression by RNA-seq, for differentials in *KMT5B/C*-deficient DIPG cells compared to wild-type.

**Supplementary Table S4** – Differentially secreted proteins in F10 and/or K5B cells compared to F8, as assessed by mass spectrometry.

**Supplementary Table S5** – Differential cis-regulated genes in F10 and/or K5B cells compared to F8, as assessed by Cicero co-accessibility analysis of single cell ATAC-seq data.

**Supplementary Table S6** – GINI index as a measure of transcriptional heterogeneity in *KMT5B/C*-deficient and wild-type DIPG cells, as applied to SmartSeq2 single cell RNA-seq.

## METHODS

### Cell culture

Patient-derived cultures HSJD-DIPG-007 and ICR-B193 were grown in stem cell media consisting of Dulbecco’s Modified Eagles Medium: Nutrient Mixture F12 (DMEM/F12), Neurobasal-A Medium, HEPES Buffer Solution 1M, sodium pyruvate solution 100nM, nonessential amino acids solution 10mM, Glutamax-I Supplement and penicillin Streptomycin solution (all Thermo Fisher, Loughborough, UK). The media was supplemented with B-27 Supplement Minus Vitamin A, (Thermo Fisher), 20ng/ml Human-EGF, 20ng/ml Human-FGF-basic-154, 20ng/ml Human-PDGF-AA, 20ng/ml Human-PDGF-BB (all Shenandoah Biotech, Warwick, PA, USA) and 2µg/ml Heparin Solution (0.2%, Stem Cell Technologies, Cambridge, UK) to constitute the complete media. In brief, cells were incubated at 37°C, 5% CO2, 95% humidity and were refed at least twice weekly with complete media. Cell authenticity was verified using short tandem repeat (STR) DNA fingerprinting.

### CRISPR/Cas9 knockdown

Parental subclone NS-F8 from HSJD-DIPG-007 was transfected with one or both plasmids expressing Cas9 as well as the following gRNAs: 5’-ATTTTGATTGATTCTGCTGA-3’/5’-CACGGGGAAGGACACCCTGA-3’ (*KMT5B*); and 5’-CAGGGGCCGTGGTGGCAACG-3’/5’-ACCAGCTGCGTGAGACCAAG-3’ (*KMT5C*) (ATUM, Newark, NJ, USA). Each plasmid was labelled with either GFP or RFP. For a clonal selection, the transfected cells were single cell flow sorted into the inner 60 wells of 96 well plates ultra-low attachment round bottom (#7007, Corning) using a Beckman Coulter MoFlo in Class IIA2 biohazard containment hood. Cells were dropped in 100µl/well of complete media supplemented with 2X growth factors, Primocin (#ant-pm, InvivoGen), Plasmocin (#ant-mpt, InvivoGen), penicillin and streptomycin (Life Technologies). The cells were sorted into 8 plates for KMT5B-KD and 5 plates for KMT5C-KD and KMT5B/C-KD, incubated at 37°C, 5% CO2 and re-fed 30µl/well twice weekly. The outer wells were filled with PBS to avoid evaporation of the media. Clones were closely monitored and once they reach over 600µm, the single sphere was broken mechanically into single cell suspension and transferred into 2 wells of a 48-well plate. The cells were expanding until maximum 200µm and collected: cells in one of the wells were pelleted down for gDNA extraction and the other one, expanded in order to be banked into StemCell Banker DMSO free (#11890, AMSbio). Edits were confirmed by sequencing after amplification using index primers based on the NexteraXT index kit V2 setA (#15052163, Illumina) and the high fidelity Taq polymerase Kapa (#KK2602, Roche). The PCR products were pooled and cleaned according to manufacturer’s instructions and quantified on an Agilent Technology 2100 Bioanalyzer using a High Sensitivity DNA chip. After denaturation, the library was diluted at 4nM and loaded on the cartridge (MiSeq reagent kit V2 Nano 500 cycles; #MS-103-1003, Illumina) at 6 pM with 15% PhiX control DNA v3 (#FC-110-3001, Illumina) and sequenced on a MiSeq System (Illumina).

### Western blot

Cell pellets were lysed by using RIPA lysis buffer (Tris 50mM pH8, NaCl 150mM, MgCl2 0.5mM, EDTA 0.5mM, Triton X-100 1%, SDS 0.5%) supplemented with protease inhibitor cocktail (Sigma, Poole, UK), phosphatase inhibitor cocktails (Sigma, Poole, UK), and sodium butyrate (Sigma, Poole, UK). Following quantification using Pierce BCA Protein Assay Kit (Thermo Fisher), equal amounts of cell extracts were loaded and membranes incubated with primary antibody overnight at 4°C, and horseradish peroxidase secondary antibody (Amersham Bioscience, Amersham, UK) for 1h at room temperature. Signal was detected with ECL Prime western blotting detection agent (Amersham Biosciences), visualised using Hyperfilm ECL (Amersham Biosciences) and analysed using an X-ray film processor in accordance with standard protocols. Primary antibodies used were H3.3K27M at 1/1000 (#ABE419 Merck Milipore), CXorf67 (EZHIP) at 1/1000 (HPA004003, Atlas Antibodies), H4K20me3 at 1/2000 (#ab9053, Abcam), H4K20me2 at 1/2000 (#ab9052, Abcam), H4K20me1 at 1/2000 (#ab9051, Abcam), H4 at 1/2000 (#ab70701, Abcam), KMT5B at 1/1000 (#LS-C161629, LSBio), KMT5C at 1/1000 (#PA5-13436, ThermoFischer), GAPDH at 1/1000 (#2118, Cell Signaling), H3K27me3 at 1/500 (#9735, Cell Signaling), H3 at 1/2000 (#9715, Cell Signaling).

### ChIP-seq

Cells were fixed with 1% formaldehyde for 15 min and quenched with 0.125 M glycine. Chromatin was isolated by the addition of lysis buffer, followed by disruption with a Dounce homogenizer. Lysates were sonicated and the DNA sheared to an average length of 300-500 bp. Genomic DNA (Input) was prepared by treating aliquots of chromatin with RNase, proteinase K and heat for de-crosslinking, followed by ethanol precipitation. Pellets were resuspended and the resulting DNA was quantified on a NanoDrop spectrophotometer. Extrapolation to the original chromatin volume allowed quantitation of the total chromatin yield. An aliquot of chromatin (30 ug) was precleared with protein A (or G; for H4K20me3) agarose beads (Invitrogen). Genomic DNA regions of interest were isolated using 4 μg of antibody against H4K16ac (#39167; 5µl), H4K20me3 (#91107; 5µl), H3K27me3 (#39155;4µl), H3K9me3 (#39161; 5µl), and H3K4me3 (#39159; 3µl). Antibody against H2Av (0.4 ug) was also present in the reaction to ensure efficient pull-down of the spike-in chromatin (Egan et al., 2016). All antibodies were from Active Motif (Carlsbad, CA, USA). Complexes were washed, eluted from the beads with SDS buffer, and subjected to RNase and proteinase K treatment. Crosslinks were reversed by incubation overnight at 65 C, and ChIP DNA was purified by phenol-chloroform extraction and ethanol precipitation. Illumina sequencing libraries were prepared from the ChIP and Input DNAs by the standard consecutive enzymatic steps of end-polishing, dA-addition, and adaptor ligation. Steps were performed on an automated system (Apollo 342, Wafergen Biosystems/Takara). After a final PCR amplification step, the resulting DNA libraries were quantified and sequenced on Illumina’s NextSeq 500 (75 nt reads, single end).

Sequences reads were aligned to the human genome hg19 using SICER (Zang et al., 2009)). PCR duplicates were removed using PICARD tool, and the read extension was counted using GAlignments. Significant peaks were identified using Model-Based Analysis for ChIP-Seq MACS version 2 (Zhang et al., 2008) using broadpeak with a p value cut-off of 10^−9^. We counted merge peaks for each mark by counting all the BAMs within the genome region. Peaks were annotated using ChIPeakAnno) with promoter region classified at peaks within +/− 2.5kb of a transcription start site (TSS). We down-sampled the tags from each mark and each sample to achieve the normalized tag counts.

### RNA-seq

RNA was isolated from cell pellets using the RNeasy Mini Kit protocol (Qiagen,74104), and measured using a Nanodrop spectrophotometer. RNA integrity was analysed and quantified using 2200-Tapestation (Agilent). A minimum of 150 ng of RNA was sequenced at Eurofins Genomics after generation of strand-specific cDNA libraries by purification of poly-A containing mRNA molecules, mRNA fragmentation, random primed cDNA synthesis (strand specific), adapter ligation and adapter specific PCR amplification. Library was then run on an Illumina Hiseq or Illumina Novaseq sequencer at 2×150 bp read length generating at least 30 million reads.

Gene expression from bulk RNASeq was quantitated in 14918 known protein-coding genes using HTSeq based upon an Ensembl gene model. Gene Set Enrichment Analysis (software.broadinstitute.org/gsea) was performed using the fgsea R package based upon pairwise comparisons. Differential expression analysis used raw HTSeq counts compared using DESeq2 and expression values were summarised both as rlog transformed expression values and RPKM.

### Drug treatment

Parental and genetically engineered cell lines were seeded at 3000 cells density per well into black wall 96-well plates (#655090, Greiner) with 50ul of complete media, 6 replicates per condition. Cells were incubated at 37C, 5% CO2. Three days after seeding, 50ul of media with either A-196 (KMT5B/C inhibitor), Talazoparib (PARP inhibitor) or DMSO were added to each well at a final concentration range between 0-10μM for A-196 and 0-1μM for Talazoparib. The cell proliferation was assessed 5 days after treatment by measuring viable cells in culture based on quantitation of the ATP present using CellTiter-Glo® Luminescent Cell Viability Assay (Promega) according to the manufacturer’s instructions using a Victor X5 Multi-label plate reader luminescence protocol (Perkin Elmer, Waltham, MA, USA). Luminescence data from each well was normalised to the median signal from DMSO-containing wells to calculate the survival fraction.

### Immunofluorescence

DIPG cells were grown on laminin pre-coated 8 well-chamber slides (Cole Palmer). Cells on chamber slides were fixed with 4% paraformaldehyde at room temperature for 15 minutes, quenched with 50mM glycine and washed with phosphate buffered saline (PBS) solution. Invasive neurospheres were collected embedded in Matrigel after 3 days of invasion assay (see method below) and were fixed in conical tube with an excess of 4% paraformaldehyde for 5h at 4°C; then washed, permeabilised with 1% TritonX-100 PBS solution for 45min. Cells were permeabilised with 0.5% Triton X-100 solution for 10 minutes at room temperature. Cells and neurospheres were then blocked with appropriate serum according to the species of secondary antibody for 1 hour at room temperature. Primary antibodies directed against 53BP1 (#CS-4937; Cell signaling) were added to the cells and incubated overnight at 4°C; H4K20me3 at 1/500 (#ab9053; Abcam), H4K20me2 at 1/1000 (#ab9052; Abcam), H4K20me1 at 1/400 (#ab9051; Abcam) were added to the neurospheres for 48h at 4C. Samples were then washed in PBS three times and incubated with Alexa Fluor488-conjugated secondary antibodies for 1 hour at room temperature for the cells or 48h at 4C for the neurospheres. Nuclei were counterstained with DAPI for all samples. The neurospheres were transferred into 8-well chamber slides. All samples were mounted with Vectashield (Vector Laboratories) and examined using a Zeiss LSM700 confocal microscope. The number of 53BP1 foci were quantified using the function Tissue Studio IF from the software Definiens Software XD64.

### Invasion

50-100μm diameter neurospheres were harvested after three days culture and invasion assays performed essentially as described (Vinci et al., 2015; Vinci et al., 2013; Vinci et al., 2018). Briefly, a total of 100μl medium was removed from ULA 96-well round-bottomed plates and 100μl of matrigel gently added to each well (6 replicates). Plates were incubated at 37°C, 5% CO2, 95% humidity for 1hr. Once the matrigel solidified, 100μl/well of culture medium was added on top. Starting from time zero, and at 3h intervals up to 6 days, automated image analysis was carried out on the live-cell imaging system IncucyteS3 (Essen Bioscience) using the *Tumor Spheroid Assays* application. Cell spread was measured after segmentation using MATLAB to perform threshold-based image segmentation on pixel intensity values followed by filtering by size, a minimum bounding circle algorithm was used to quantify cell spread from the resulting segmentation masks (n>=3).

### Secretome

Established patient-derived cells were seeded and cultured for 5 days in 3D when the neurospheres reach 50-100μm diameter. The cells were then washed with HBSS and gently seeded in media deprived with B-27 for 24h at 37°C, 5% CO2, 95% humidity. The secretomes were harvested, filtered through 0.22um filters (MillexGP, Sigma) and concentrated in Amicon® Ultra Centrifugal Filters, 3kDa cut-off (Millipore). The concentrated secretome was then analysed by Mass Spectrometry. 100 µg of secretome proteins per sample was reduced by TCEP (Tris(2-carboxyethyl)phosphine, Sigma), alkylated by iodoacetamide (Sigma). Proteins were precipitated by 20% (w/v) trichloroacetic acid to remove the amino acids in the samples. Proteins were resuspended in 100 mM TEAB buffer (triethylammonium bicarbonate, Sigma) and then digested by trypsin (Pierce MS grade, Thermo Fisher) for 18 hours at 37°C. Peptides were labelled by 0.8 mg of TMT10plex according to the manufacturer’s instruction.

Samples were pooled and dried in SpeedVac, and then fractionated either by Pierce™ high pH reversed-phase peptide fractionation spin column to 8 fractions from 10% - 50% ACN/0.1% triethlyamine as instructed by the manufacturer, or online on an XBridge BEH C18 column (2.1 mm i.d. x 150 mm, Waters) to 6 concatenated fractions from a 30 min linear gradient of 5% - 35% ACN/0.1% NH_4_OH and run time at 60 min, where fractions were collected at every 30 s from 5 – 50 min. Fractionated peptides were dried in SpeedVac, then reconstituted in 0.5% formic acid (FA)/H_2_O and injected for on-line LC-MS/MS analysis on an Orbitraip Fusion Lumos hybrid mass spectrometer coupled with an Ultimate 3000 RSLCnano UPLC system (both from Thermo Fisher). Peptides were first loaded and desalted on a PepMap C18 nano trap (100 µm i.d. x 20 mm, 100 Å, 5µ) then peptides were separated on a PepMap C18 column (75 µm i.d. x 500 mm, 2 µm) at a flow rate at 300 nl/min. The LC gradient time was either 90 min from 10% - 38%B (run time 120 min), or 180 min from 5% - 32%B (run time 210 min), where B was 80%ACN/0.1%FA. The MS acquisition used MS3 level quantification with Synchronous Precursor Selection (SPS) with the Top Speed 3s cycle time. Briefly, the Orbitrap full MS survey scan was m/z 375 – 1500 with the resolution 120,000 at m/z 200, with AGC set at 400,000 and 50 ms maximum injection time. Multiply charged ions (z = 2 – 5) with intensity threshold at 5000 were fragmented in ion trap at 35% collision energy, with AGC at 10,000 and 35 ms maximum injection time, and isolation width at 0.7 Da in quadrupole. The top 5 MS2 fragment ions were SPS selected with the isolation width at 0.7 Da, and fragmented in HCD at 65% collision energy, and detected in the Orbitrap to get the report ions’ intensities at a better accuracy. The resolution was set at 50,000, and the AGC at 50,000 with maximum injection time at 86 ms. The dynamic exclusion was set 40 s with ± 10 ppm exclusion window.

The raw files were processed with Proteome Discoverer 2.3 or 2.4 (Thermo Fisher) using the Sequest HT search engine. Spectra were searched against fasta files of reviewed *Uniprot homo Sapiens* entries (April 2019 for or February 2020) and an in-house contaminate database. Search parameters were: trypsin with 2 maximum miss-cleavage sites, mass tolerances at 20 ppm for Precursor, and 0.5 Da for fragment ions, dynamic modifications of Deamidated (N, Q), Oxidation (M) and Acetyl (Protein N-terminius), static modifications of Carbamidomethyl (C) and TMT6plex (Peptide N-terminus and K). Peptides were validated by Percolator with q-value set at 0.01 (strict) and 0.05 (relaxed). The TMT6plex reporter ion quantifier included 20 ppm integration tolerance on the most confident centroid peak at the MS3 level. Only unique peptides were used for quantification. The co-Isolation threshold was set at 100%. Peptides with average reported S/N>3 were used for protein quantification, and the SPS mass matches threshold was set at 55%. The abundance normalization mode used the total peptide amount, and scaling mode used on all average. Only master proteins were reported.

### Single cell RNA-Seq

Each sample was prepared as a single cell suspension, with neurospheres grown until ∼100µm diameter, pelleted down, gently dissociated with accutase (#A6964, Sigma), washed and resuspended with complete media, counted and then washed again in PBS with 0.04% BSA. Single cell suspensions were processed through the 10X Chromium Single Cell Platform using Chromium Single Cell 3’ v3**.1** Chemistry (10X Genomics, Pleasanton, CA) following the manufacturer’s protocol. A total of 4,000 cells were added to each channel of a chip to be partitioned into Gel Beads in Emulsion (GEMs) in the Chromium instrument, followed by cell lysis and barcoded reverse transcription of RNA in the droplets. Breaking of the emulsion was followed by amplification, fragmentation, and addition of adaptor and sample index. Single Cells 3’ Gene Expression libraries were sequenced on the NextSeq 500 and processed with Cell Ranger analysis pipeline. For SmartSeq2, Single cell suspensions were sorted for viable cells (calcein positive,TO-PRO-3 negative) into 96-well plates for whole transcriptome amplification, library preparation and pooling for sequencing on a NextSeq 500 as previously described (Picelli et al., 2014). Raw sequencing reads were aligned to hg19 genome using Bowtie, and gene expression was quantified as transcripts per million (TPM) with RSEM. Analysis was performed using Seurat version 3.0 (Butler et al., 2018)) and visualised using UMAP projection. Copy number profiles from single cell RNA sequencing were generated with infercnv (Patel et al., 2014).

### Single cell ATAC-Seq

Each sample was prepared as a single cell suspension, with neurospheres grown until **∼**100µm diameter, pelleted down, gently dissociated with accutase (#A6964, Sigma), washed and resuspended with cryopreservation media (STEM-CELLBANKER; Amsbio) at 4 million cells/mL, then cells were cryopreserved in a Mr Frosty. Cells were thawed in a 37°C water bath and prepared as described by 10X Genomics Demonstrated Protocol – Nuclei Isolation for Single Cell ATAC Sequencing Rev B. Briefly, cell pellets were resuspended in lysis buffer and incubated on ice for 5 minutes. Lysed cells were washed, strained, and nuclei were resuspended and counted using a Countess II FL Automated Cell Counter. Isolated nuclei were then used as input following the 10X Genomics Chromium Next GEM Single Cell ATAC Reagent Kits v1.1 manual. Targeting a 5,000 nuclei recovery, samples were added to the tagmentation reaction, loaded into the Chromium Controller for nuclei barcoding, and prepared for library construction following manufacturer’s protocol (10X Genomics PN-1000175). Resulting libraries were quantified using the KAPA Library Quantification Kit for Illumina platforms (KAPA Biosystems), and sequenced with PE34 sequencing on the NextSeq 500/550 sequencer (Illumina).

Sequenced data were processed with the Cell Ranger ATAC software, with alignment to the human (hg38) genome. The Cell Ranger output files were used as input to Active Motif’s proprietary analysis program, which creates Excel tables containing detailed information on cluster-specific peak locations, gene annotations, and motif enrichment. The alignment files generated by Cell Ranger were also processed as pseudo-bulk ATAC-Seq samples. Duplicate reads were removed, only reads mapping as matched pairs and only uniquely mapped reads (mapping quality >= 1) were used for further analysis. Alignments were extended in silico at their 3’-ends to a length of 200 bp and assigned to 32-nt bins along the genome. The resulting histograms (genomic “signal maps”) were stored in bigWig files. Peaks were identified using the MACS 2.1.0 algorithm at a cutoff of p-value 1×10-7, without control file, and with the – nomodel option. Peaks that were on the ENCODE blacklist of known false ChIP-Seq peaks were removed. Signal maps and peak locations were used as input data to Active Motif’s proprietary analysis program, which creates Excel tables containing detailed information on sample comparison, peak metrics, peak locations and gene annotations.

### Statistics

Statistical analysis was carried out using R 3.5.0 (www.r-project.org) and GraphPad Prism 8. Categorical comparisons of counts were carried out using Fishers exact test, comparisons between groups of continuous variables employed Student’s t-test or ANOVA. Univariate differences in survival were analysed by the Kaplan-Meier method and significance determined by the log-rank test. All tests were two-sided and a p value of less than 0.05 was considered significant. Normalised and scaled abundances proteins were compared by using a Student’s t-test adjusted for false discovery rate (FDR) according to Benjamini and Hochberg. For gene expression data analysis multiple testing corrections were made using FDR according to Benjamini and Hochberg. Effects of drug treatment on survival as the primary endpoint and overall survival in the orthotopic *in vivo* models were assessed using Mantel Cox log-rank test. Adjusted P-values < 0.05 was considered significant.

### Data accessibility

All newly generated sequencing data have been deposited in the European Genome-phenome Archive (ebi.ac.uk/ega) with accession number EGAS00001005365.

## Notes

*Conflict of interest statement:* The authors declare no conflicts of interest pertaining to this manuscript.

### Competing Interest Statement

The authors have declared no competing interest.

